# Epigenetic mechanisms regulating CD8+ T cell senescence in aging humans

**DOI:** 10.1101/2025.01.17.633634

**Authors:** Paolo S. Turano, Elizabeth Akbulut, Hannah K. Dewald, Themistoklis Vasilopoulos, Patricia Fitzgerald-Bocarsly, Utz Herbig, Ricardo Iván Martínez-Zamudio

## Abstract

Aging leads to the decline of immunity, rendering the elderly susceptible to infection and disease. In the CD8+ T cell compartment, aging leads to a substantial increase of cells with high levels of senescence-associated ß-galactosidase activity (SA-ßGal) and other senescence characteristics, including a pro-inflammatory transcriptome and impaired proliferative potential. Using senescent cell isolation coupled with multiomic profiling, here we characterized the epigenetic mechanisms regulating CD8+ T cell senescence in a cohort of younger and older donors. High levels of SA-ßGal activity defined changes to global transcriptomes and chromatin accessibility landscapes, with a minor effect of age. Widespread enhancer remodeling was required for the repression of functional CD8+ T cell genes and upregulation of inflammatory and secretory pathway genes. Mechanistically, the senescence program in CD8+ T cells was controlled by chromatin state-specific transcription factor (TF) networks whose composition was largely insensitive to donor age. Pharmacological inhibition of TF network nodes AP1, KLF5, and RUNX2 modulated the transcriptional output, demonstrating the feasibility of TF network perturbation as an approach to modulate CD8+ T cell senescence. Further, CD8+ T cell senescence gene signatures faithfully predicted refractoriness to chimeric antigen receptor (CAR) T-cell therapy in a cohort of diffuse large B cell lymphomas and were highly enriched in the transcriptomes of peripheral CD8+ T cells of individuals with active systemic lupus erythematosus. Collectively, our findings demonstrate the potential of multiomic profiling in identifying key regulators of senescence across cell types and suggest a critical role of senescent CD8+ T cells in disease progression.

## INTRODUCTION

Aging leads to increased susceptibility to infection and disease in older populations. Recent evidence has established cellular senescence, a dysfunctional cell state defined by a stable proliferative arrest and a pleiotropic secretome (the senescence-associated secretory phenotype; [SASP]) that alters the tissue microenvironment, as a critical contributor to the aging process ^1,2^. Stresses that perturb cell identity activate the senescence program to signal the presence of tissue damage and coordinate subsequent repair ^3,4^. Through the SASP, senescent cells recruit innate and adaptive cell types of the immune system for their removal, limiting the deleterious effects caused by the prolonged stay of senescent cells in tissues ^5,6^. However, the capacity of the immune system to remove senescent cells from tissues decreases with age. This overall decline in immune function, known as immunosenescence, is defined by impaired thymopoiesis, disrupted CD4+ to CD8+ T cell ratios and chronic low-level inflammation, and contributes to the increased morbidity and mortality in older adults ^7,8^. Whether the immune system itself undergoes senescence with age remains poorly understood. However, defining the molecular mechanisms that drive senescence of the immune system could reveal opportunities to developing therapeutic interventions to restore immune function in the older population to increase healthspan.

In the CD8+ T cell compartment, the age-associated decline in immune function is linked to the expansion of highly differentiated subsets that acquire dysfunctional markers, including PD1 (PDCD1), KLRG1, and CD57 ^9–13^, decreased proliferative capacity upon stimulation and nonspecific cytotoxic capacity through upregulation of NKG2D ^14^. Using a senescent cell isolation method based on the detection of senescence-associated beta-galactosidase (SA-ßGal) activity, we recently revealed a drastic increase in CD8+ T cells with features of senescence, including high levels of cell cycle inhibitors CDKN1A (p21) and CDKN2A (p16), an inflammatory SASP and telomeric DNA damage, across all differentiation states and encompassing several described CD8+ T cell dysfunction markers in the peripheral blood of older individuals ^15^, suggesting that the transition to senescence plays an outsized role in the dysfunction of CD8+ T cells with age. Despite their proliferative arrest, recent epigenetic studies show that senescent cells are dynamic entities that continuously evolve ^16–18^. This nature of the senescence state underlies its heterogeneity but also exposes liabilities in senescent cells that can potentially facilitate their reprogramming to a functional identity or promote their elimination ^19,20^. Although recent work on the epigenetic regulation of CD8+ T cells during aging has established a correlation between dedicated usage of transcription factors (TFs) to mediate differentiation, aging, and dysfunctional state-specific gene expression programs ^21–24^, the gene regulatory mechanisms that mediate the transition to senescence in CD8+ T cells during aging in humans are not known.

In this study, we combined senescent cell isolation with multiomic profiling to define the gene regulatory mechanisms driving CD8+ T cell senescence in a cohort of younger and older donors. This approach identified the acquisition of senescence as the major driver of epigenome and gene expression changes in CD8+ T cells, with a minor contribution of age. We establish enhancer remodeling as a key mechanism driving senescence-associated gene expression in CD8+ T cells, define and modulate the TF networks controlling the senescence program, and develop CD8+ T cell senescence gene signatures with diagnostic and prognostic potential for CAR-T cell therapy and systemic lupus erythematosus (SLE).

## RESULTS

### Transcriptional landscape of senescent CD8+ T cells

We leveraged our approach to detect and isolate live senescent cells and purified live non-senescent (low SA-ßGal) and senescent (high SA-ßGal) CD8+ T cells from peripheral blood of a cohort of 14 healthy donors (7 young donors in their 20-27 years old and 7 older donors 55 years or older) and performed bulk RNA-sequencing. Consistent with our previous results ^15^, principal component analysis (PCA) and hierarchical clustering confirmed that the transcriptome of CD8+ T cells from young and older donors partitioned based mostly on SA-ßGal content (up to 69% of the variance) with a small contribution of donor variability (up to 9% of the variance) and a negligible contribution of age (**Figures 1A-B**). Further characterization of their transcriptomes utilizing a self-organizing map (SOM) algorithm ^25^ revealed generally consistent expression portraits of non-senescent and senescent CD8+ T cells, although donor-specific differences were also detected (**Figures 1C** and **Supplementary** Figure 1A). For instance, genes in clusters K and M, which enriched for inflammatory pathways, exhibited consistent expression levels across age and senescence status (**Supplementary** Figures 1B-D). In contrast, small but detectable age and senescence status-specific levels were detected for EMT- and coagulation-associated genes in clusters A and H (**Supplementary** Figures 1B-D). Thus, non-senescent and senescent CD8+ T cells exhibit coherent expression portraits across donors independently of age. We then performed differential expression analysis and identified 4,703 differentially expressed genes (DEGs), which split into two modules characterized by non-senescent (blue) and senescent (turquoise) CD8+ T cells irrespective of age (**Figures 1D-E**). Pathway analysis revealed extensive representation and diversity of senescence-associated pathways on the SA-ßGal-high-specific turquoise module, including cell cycle regulation, P53 pathway and multiple inflammatory signaling and secretory pathways. In contrast, the blue module characterizing non-senescent CD8+ T cells was highly specialized, with enrichment of proliferation-associated MYC targets (**Figure 1F**). We identified a small number of age-specific DEGs in non-senescent (127) and senescent (62) CD8+ T cells whose expression pattern exhibited more pronounced donor-specific differences than senescence-specific DEGs (**Figure 1G** and **Supplementary** Figures 1E-F). These genes were overrepresented in a small subset of pathways, including EMT, hypoxia, and glycolysis, indicating that age-specific changes to CD8+ T cell expression are specialized. In contrast, senescence-specific DEGs exhibited substantial diversity of biological function (**Figure 1H**). Together, our analysis of transcriptomes of CD8+ T cells from young and older donors highlights the acquisition of senescence as the major driver of gene expression changes with age.

**Figure 1.**
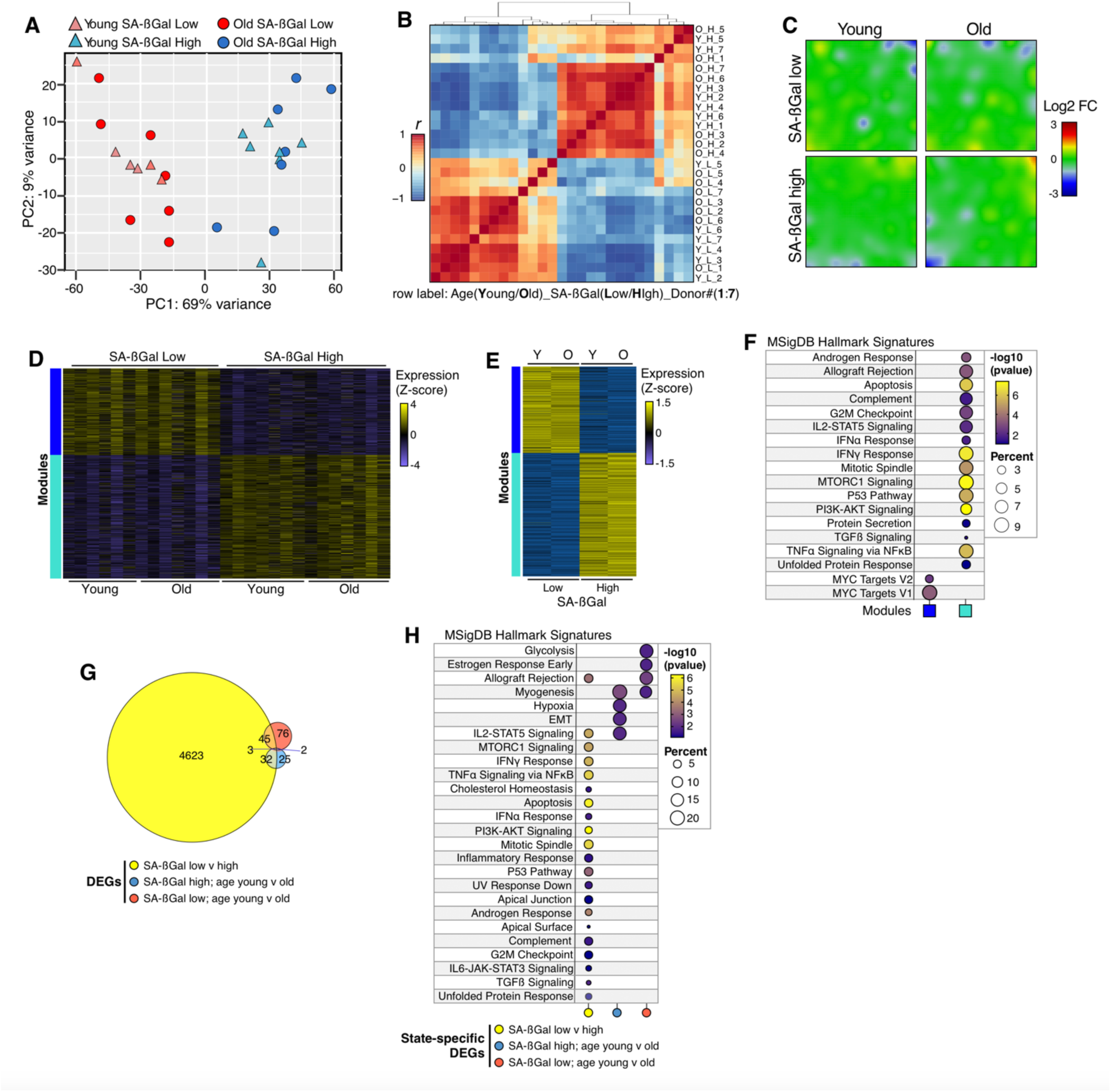
Transcriptional landscape of senescent CD8+ T cells. A. Principal component analysis (PCA) of transcriptomes (reads on exons) of SA-ßGal-low (red) and SA-ßGal-high (blue) CD8+ T cells isolated from younger (triangles) and older (circles) individuals. **B.** Heatmap showing the correlation (r) between the CD8+ T cell transcriptomes of the indicated donors. **C.** Averaged self-organizing maps (SOMs) of transcriptomes of SA-ßGal-low and SA-ßGal-high CD8+ T cells from younger and older individuals. Individual SOMs are shown in Figure S1A. Expression is represented as log_2_ fold change. **D, E.** Individual and averaged heatmaps of color-coded modules of differentially expressed genes (DEGs) in SA-ßGal-low and SA-ßGal-high CD8+ T cells from younger and older donors (row Z-score; n = 1941 genes for the blue module; n = 2762 genes for the turquoise module). **F.** Functional overrepresentation analysis map showing significant associations of the MSigDB hallmark gene sets for each expression module described in E. Circle fill is color coded according to the false discovery rate (FDR)-corrected p value from a hypergeometric distribution test. Circle size is proportional to the percentage of genes in each MSigDB gene set. **G.** Intersections of DEGs between the indicated comparisons (bottom). **H.** As in F but for the DEG sets described in G. A-H were generated from transcriptomic data from SA-ßGal-low and SA-ßGal-high CD8+ T cells of seven younger and seven older donors (A-H).

### SA-ßGal activity levels define the accessible chromatin of CD8+ T cells

To gain insights into the epigenetic mechanisms driving CD8+ T cell senescence, we performed the assay for transposase accessible chromatin followed by sequencing (ATAC-seq) ^26^ on sorted SA-ßGal-low and SA-ßgal-high CD8+ T cells from 4 younger and 4 older donors. Consistent with the results from our transcriptome analysis, the accessible chromatin of CD8+ T cells was defined mostly by SA-ßGal levels, with a small contribution of inter-individual variation and a negligible effect of age, as shown by PCA of total reads on peaks (**Figure 2A**). We identified 6,936 differentially accessible regions (DARs) which partitioned into two distinct modules, I and II, that defined non-senescent and senescent CD8+ T cells. The patterns of DARs were generally consistent across individuals, although we noticed a reduction of signal intensity on the averaged DARs of both SA-ßGal-low and SA-ßGal-high CD8+ T cells of older donors (**Figures 2B-C**). Annotation and integration of DARs with gene expression retrieved 1,510 DEGs (549 specific to module I, 814 to module II, 147 in common) linked to the nearest DARs (**Figure 2D**), accounting for ∼32% of the DEGs in senescent CD8+ T cells, suggesting the existence of additional regulatory layers to the transcriptional output. Most DARs from both modules localized to intronic and intergenic regions within 40 kilobases from the transcription start site (TSS) of DEGs (**Figures 2E-F** and **Supplementary** Figures 2A-B). Most DEGs were primarily linked to a single DAR (**Figures 2G** and **Supplementary** Figure 2C), indicating that a single regulatory chromatin binding event is sufficient to drive changes to their expression ^27^. The directionality and magnitude of chromatin accessibility changes generally reflected those of their linked DEGs, consistent with previous reports ^16,27^. For instance, decreased accessibility of DARs in module II in senescent CD8+ T cells correlated with repression of metabolism-related DEGs characteristic of SA-ßGal-low CD8+ T cells (**Supplementary** Figures 2D-E). In contrast, most DEGs linked to open DARs in module I in senescent CD8+ T cells were highly expressed and represented senescence pathways including the secretory phenotype (SASP) and cell cycle control (**Figures 2H-I**). Collectively, our results demonstrate that the acquisition of senescence is the major driver of chromatin accessibility changes in CD8+ T cells.

**Figure 2.**
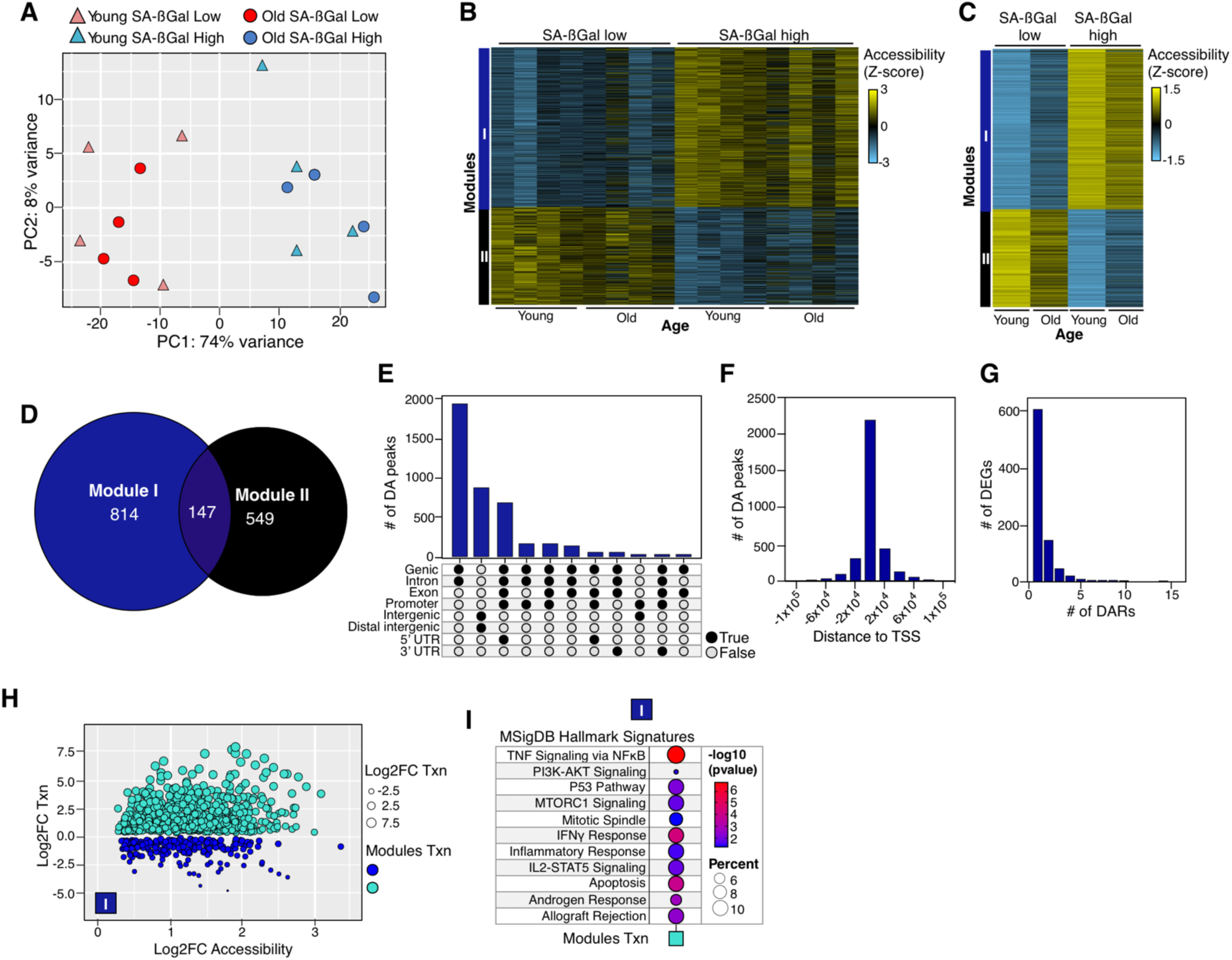
Senescence and not age drives changes in chromatin accessibility of CD8+ T cells. A. Principal component analysis (PCA) of chromatin accessibility (reads on peaks) of SA-ßGal-low (red) and SA-ßGal-high (blue) CD8+ T cells isolated from younger (triangles) and older (circles) individuals. **B, C.** Individual and averaged color-coded modules of differentially accessible (DA) peaks in SA-ßGal-low and SA-ßGal-high CD8+ T cells from younger and older donors (n = 2636 peaks for the black module; n = 4300 genes for the dark blue module). **D.** Intersections of senescence-specific DEGs linked to each DA peak module identified in B. **E.** UpSet plot showing the frequency distribution of opening DA peaks (dark blue module) with the indicated genomic features. **F.** Frequency distribution showing the occurrence of opening DA peaks relative to the TSS of the nearest annotated gene. **G.** Histogram plot showing the number of DEGs linked to opening DA peaks. **H.** Correlation between chromatin accessibility and expression of DEGs linked to opening DA peaks. Individual DEGs from the blue and turquoise modules from 1C are shown as color coded circles. Circle size is proportional to the log2 fold change in expression. **I.** Functional overrepresentation analysis map showing significant associations of the MSigDB hallmark gene sets for the indicated DEGs linked to opening DA peaks. Circle fill is color coded according to the false discovery rate (FDR)-corrected p value from a hypergeometric distribution test. Circle size is proportional to the percentage of genes in each MSigDB gene set. Chromatin accessibility data (A-H) were generated from SA-ßGal-low and SA-ßGal-high CD8+ T cells from four younger donors and four older donors and integrated with the transcriptomic data from SA-ßGal-low and SA-ßGal-high CD8+ T cells of seven younger and seven older donors (G-I).

### Enhancer remodeling dictates the transcriptional output of senescent CD8+ T cells

Previous studies have shown a critical role of enhancers in the timely execution of senescence programs across distinct cell types ^17,28,29^. We therefore determined the enhancer dynamics of SA-ßGal-low and SA-ßGal-high CD8+ T cells. To this end, we performed cleavage under targets and tagmentation (CUT&Tag) ^30^ to map poised enhancers (H3K4me1-defined regions), Polycomb repressed chromatin (H3K27me3-defined regions) and active chromatin (H3K27ac-defined regions) genome-wide. We subsequently combined histone modification and chromatin accessibility datasets and performed a chromatin state transition analysis ^31^, modeling the acquisition of senescence as a transition from SA-ßGal-low to SA-ßGal-high state. Consistent with our published results of fibroblasts undergoing oncogene-induced senescence (OIS) ^16,32^, the mappable genome of SA-ßGal-low and SA-ßGal-high CD8+ T cells was largely (<80%) devoid of the epigenetic features profiled, with only ∼16% of the genome attributed to these (**Supplementary** Figure 3A). Intriguingly, and in contrast to our previous results on fibroblasts, inactivation of enhancers to the unmarked, poised and Polycomb repressed states, followed by enhancer activation from the unmarked state, characterized senescent CD8+ T cells (**Figures 3A-B** and **Supplementary** Figure 3B). Accordingly, enhancer inactivation was linked to the reduced transcriptional output of nearby genes and was significantly correlated to the repression of SA-ßGal-low-specific DEGs while the activation of enhancers robustly correlated with increased expression of senescence-specific DEGs in SA-ßGal-high CD8+ T cells (**Figures 3C** and **Supplementary** Figure 3C). For instance, induction of *TBKBP1*, an adaptor protein to pro-inflammatory kinase TBK1 ^33^, and *TBX21*, a transcription factor involved in regulating exhaustion ^34^ and senescence (see below), is linked to enhancer activation from the poised and Polycomb repressed states, respectively (**Figure 3D**). In contrast, repression of *LAYN*, an activator of integrin function and mediator of antitumor immunity ^35^, is associated with H3K27me3 deposition across its entire gene body and highlights the possibility of reduced antitumor surveillance capacity of senescent CD8+ T cells. Collectively, our comprehensive chromatin state analysis highlights local enhancer remodeling as a critical epigenetic mechanism driving the senescence gene expression program in CD8+ T cells.

**Figure 3.**
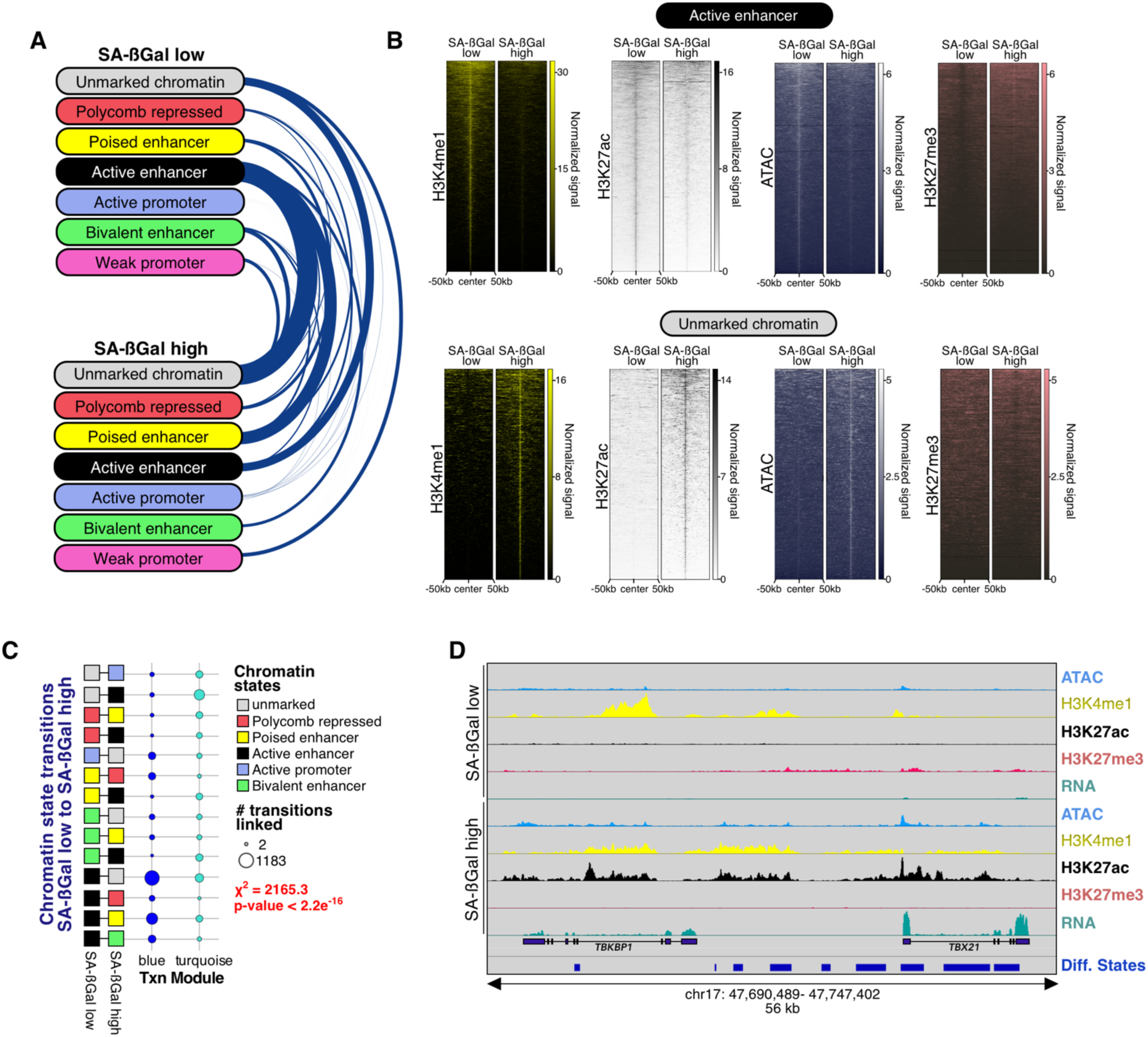
Enhancer remodeling drives the transition to senescence in CD8+ T cells. A. Arc-plot representation summarizing the evolution of the most prominent chromatin states as CD8+ T cells transition into senescence. The edge width is proportional to the number of 200 bp bins undergoing the indicated chromatin state transition. **B.** Genome-wide signal intensity evolution heatmaps of H3K4me1, H3K27ac, ATAC-seq and H3K27me3 at active enhancers (top) and unmarked chromatin (bottom) in SA-ßGal-low and SA-ßGal-high CD8+ T cells. **C.** Dot-plot of the correspondence analysis linking the indicated chromatin state transitions to gene expression modules. The size of the circles is proportional to the number of 200 bp bins undergoing the indicated transitions linked to the gene expression modules. The significance of the association was calculated using a chi-squared test and the p-value is shown. **D.** Representative genome browser snapshot of the *TBKBP1* and *TBX21* loci showing signal tracks for ATAC-seq, H3K4me1, H3K27ac and H3K27me3 in SA-ßGal-low and SA-ßGal-high CD8+ T cells. Cut&Tag data (A-D) were generated from SA-ßGal-low and SA-ßGal-high CD8+ T cells from three younger donors and integrated with the transcriptomic data from SA-ßGal-low and SA-ßGal-high CD8+ T cells of seven younger and seven older donors (C, D).

### Local activation of the transcription factor network underpins the transition to senescence in CD8+ T cells

Transcription factor (TF) networks orchestrate cell fate transitions by modulating chromatin structure and controlling the timing and magnitude of the gene expression output ^36,37^. To define the function and dynamics of the TF networks that mediate the senescence program in CD8+ T cells, we searched for TFs exhibiting differential expression between SA-ßGal-low and SA-ßGal-high CD8+ T cells from younger and older donor. Surprisingly, we identified 455 differentially regulated TFs, representing ∼40% of all detectable TFs in CD8+ T cells, which partitioned essentially based on SA-ßGal level and irrespective of age (**Figure 4A**). The high number of differentially regulated genes between SA-ßGal-low and SA-ßGal-high CD8+ T cells suggested a large rearrangement to their gene regulatory network as these cells transitioned to senescence. To address this possibility, we quantified TF binding instances on DAR modules (see **Figure 2**) of SA-ßGal-low and SA-ßGal-high CD8+ T cells of younger and older donors. The KLF family of TFs emerged as the most pervasively bound TFs to closing and opening chromatin in both age groups, irrespective of senescence status, suggesting a pioneer role for KLF TFs in CD8+ T cells ^38^ (**Figures 4B-E** and **Supplementary Figures S4A-D**). Interestingly, members of the AP1 family of TFs, were prebound to senescence-specific open chromatin irrespective of senescence status, indicating that, along KLFs and additional TFs, AP1 TFs act as bookmarking agents for the timely execution of the senescence program ^17^ in CD8+ T cells. We subsequently integrated differential TF binding activity at opening and closing chromatin with their respective transcriptional dynamics, which revealed a highly dynamic TF network that relied on both constitutively as well as differentially expressed TFs (**Figure 4F** and **Supplementary** Figure 4E). For instance, binding activity of AP1 (FOSL2, FOS) and AP1-associated (MAFs, and BACHs) TFs, as well as TBX1 and EOMES, was highest in the open chromatin in SA-ßGal-high CD8+ T cells (**Figure 4F**, top insets) while the binding of proliferation-associated TFs, E2F6, EGR2 and MYC was reduced (**Figure 4F**, bottom insets). Similarly, consistently higher activity of KLFs, SP NFY and REL TFs was observed at closing chromatin of senescent CD8+ T cells (**Supplementary** Figure 4E, bottom insets). Mapping of the TF co-binding interactions revealed specific TF networks at opening and closing chromatin of senescent CD8+ T cells. AP1, KLF and SP TFs mediated interactions with most other TFs at the open chromatin of SA-ßGal-high CD8+ T cells in both younger and older donors (**Figures 4G-H** and see below), while KLF TFs essentially dominated the TF interactome, including interactions with cell cycle and chromatin architectural TFs MYC, MAX and CTCFL, at closing chromatin (**Supplementary Figures S4F-G**). To visualize the structure and dynamicity of the TF interactome, we constructed TF network models by leveraging the co-binding maps and expression status of TFs at opening and closing chromatin. The CD8+ T cell TF networks were composed of both constitutively and differentially expressed TFs and displayed a hierarchical structure composed of top, core, and bottom layers. As the co-binding interactions flowed from top to bottom, there was an increasing number of nodes with increasing dynamicity, which bound progressively less regions (**Figure 5** and **Supplemental Figure 5**). Interestingly, while the structure of the TF networks was remarkably conserved in younger and older donors, these networks displayed unique compositions based on their association with opening or closing chromatin in CD8+ T cells. The opening chromatin TF network exhibited a rather complex top layer composed of 15 nodes, with AP1, RUNX, ETS, and TBX TFs occupying most regions and connecting to most nodes in the core and bottom layers (**Figure 5**), while KLF TFs controlled most interactions with the remaining nodes in the network at closing chromatin (**Supplemental Figure 5**). Interestingly, a node composed of KLF, SP, AP1, and TCF TFs played a critical role in open chromatin by acting as a relay point between the top and bottom layers (**Figure 5**), highlighting the versatility of TF node usage depending on the chromatin context ^39,40^. Although these networks display some age-specific differences in TF usage and node composition of the core and bottom (see **Figure 4F** mid inset, **Figure 5** and **Supplemental Figure 5** compare core and bottom layers), our comprehensive analyses of TF binding demonstrate that CD8+ T cells of both younger and older individuals rely on the activation of differentially accessible chromatin-specific TF networks of remarkably similar composition to implement senescence-associated gene expression.

**Figure 4.**
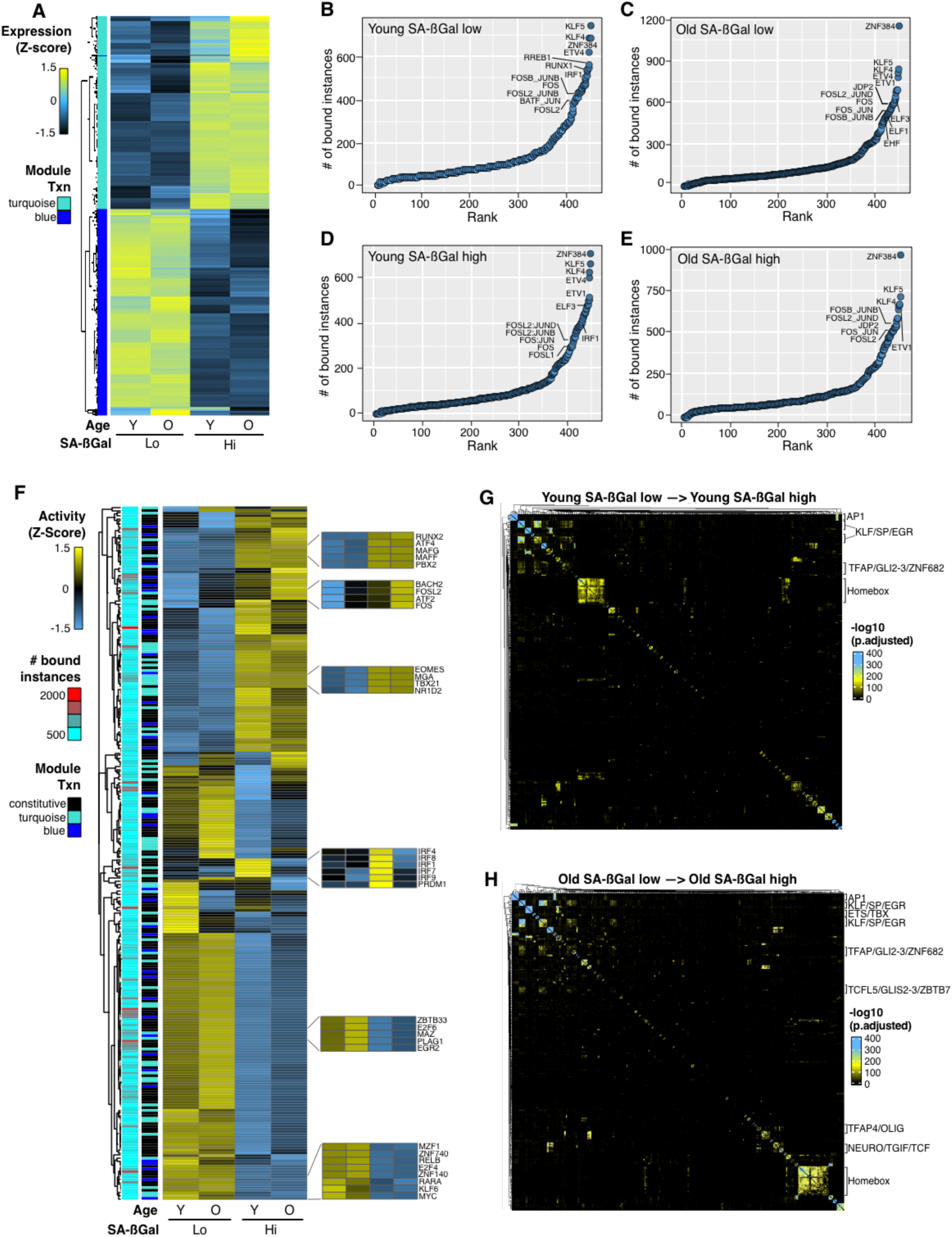
Transcription factor network dynamics of senescent CD8+ T cells. A. Averaged expression heatmaps of differentially expressed TFs in the corresponding DEG modules of SA-ßGal-low and SA-ßGal-high CD8+ T cells from younger and older individuals. **B, C, D, E.** Rank plots showing the summed binding instances per TF in the senescence-specific open chromatin of SA-ßGal-low (B, C) and SA-ßGal-high (D, E) of CD8+ T cells of younger and older donors. The TFs with the most binding instances are labelled. **F.** Heatmap of differential TF chromatin binding activity (row Z-score) at senescence-specific opening chromatin of SA-ßGal-low and SA-ßGal-high CD8+ T cells from younger and older donors. The left annotation heatmaps show the number of bound instances per TF (cyan to red) and their gene expression (TXN) category (i.e., constitutively [black] or differentially expressed according to the module color shown in Figures 1D and E. Insets: chromatin binding activity of age- and cell state-specific TFs. **G, H.** TF co-binding interaction matrices in the senescence-specific open chromatin of CD8+ T cells from younger (G) and older (H) donors transitioning into senescence. All binding instances across the two cell states (non-senescent and senescent) are collapsed onto the matrices and clustered using Ward’s aggregation criterion. The corresponding q values of the interactions are projected onto the clustering and represented in a color scale defined by their significance using a hypergeometric distribution test. A was generated from averaged RNA-seq data sets from SA-ßGal-low and SA-ßGal-high CD8+ T cells from seven younger and seven older donors. B-H were generated from pooled ATAC-seq datasets from SA-ßGal-low and SA-ßGal-high CD8+ T cells from four younger and four older donors.

**Figure 5.**
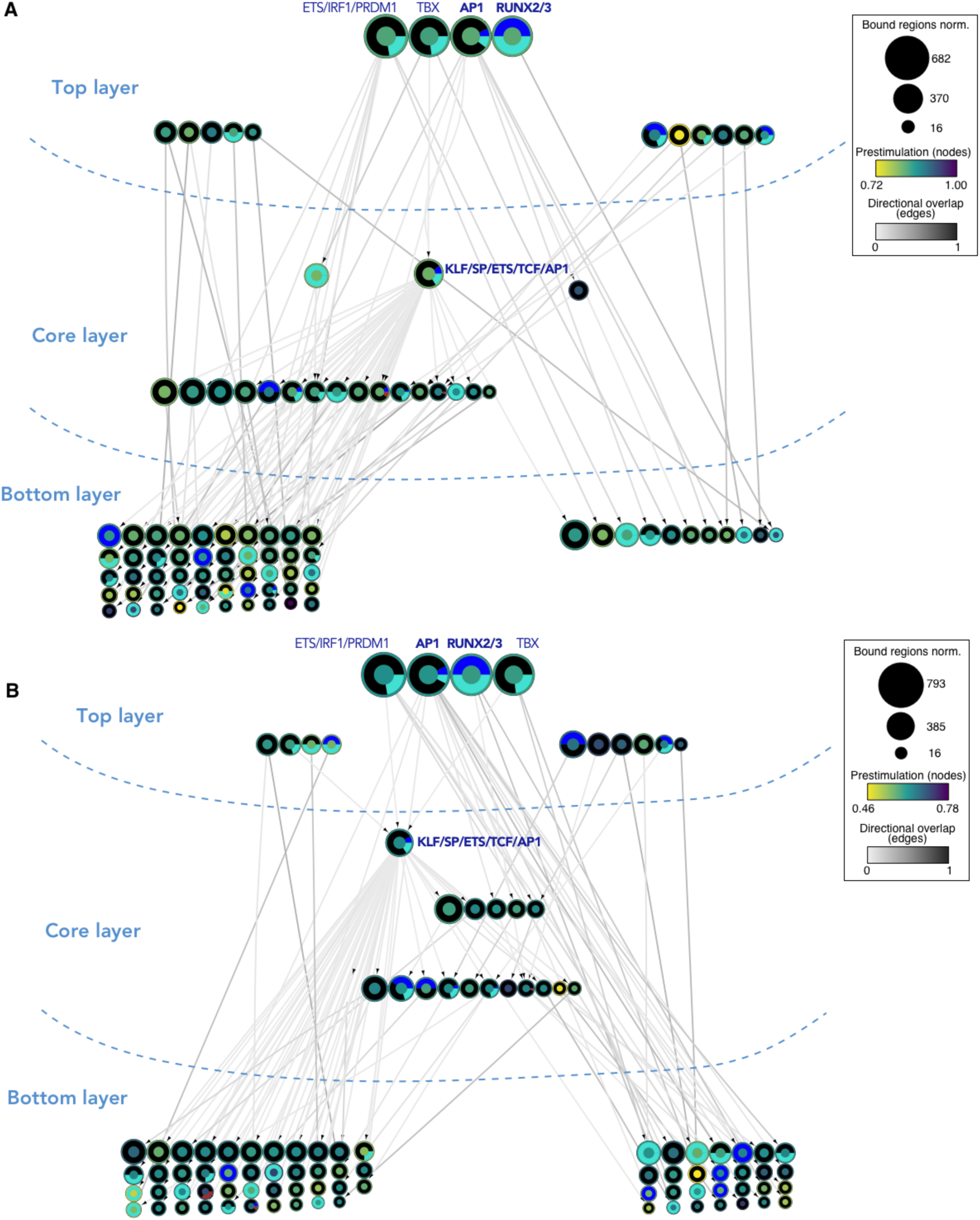
Distinct hierarchical TF networks control the senescence program of CD8+ T cells. A, B. TF networks for the senescence-specific open chromatin module (Figure 2B, dark blue module) in CD8+ T cells from younger (A) and older (B) donors transitioning into senescence. TFs (nodes) are represented as circles. Oriented edges (arrows) connecting nodes indicate that at least 15% of the regions bound by a given TF in the bottom and core layers were bound by the interacting TF in the core and top layers, respectively, in SA-ßGal-low CD8+ T cells. Nodes represent strongly connected components (SCCs) (i.e., regulons of multiple TF co-binding interactions) to facilitate visualization. The fill color of the node’s inner circle is based on the normalized dynamicity (prestimulation) of TFs. The fill color of the outer ring indicates whether the TF is constitutively expressed (black) or belongs to a transcriptomic module as described in Figure 1D. The node’s size is proportional to the bound regions by a given TF(s). Each network has three layers: (1) the top layer with no incoming edges, (2) the core layer with incoming and outgoing edges, and (3) the bottom layer with no outgoing edges. Top layer most prominent TFs and the highly connected KLF/SP SCC are labelled. Bold labels indicate TFs targeted with pharmacological inhibitors (see Figures 6 and S6). Networks were generated from pooled ATAC-seq datasets from SA-ßGal-low and SA-ßGal-high CD8+ T cells from four younger and four older donors.

### Rational targeting of the TF network to modulate senescence-associated gene expression in CD8+ T cells

The hierarchical structure of TF networks provides an opportunity for their targeted perturbation to manipulate cell states ^16^. Using this logic, we purified SA-ßGal-low and -high CD8+ T cells, subsequently cultured them in the presence of IL2 for three days and determined their transcriptomic profiles. We identified 4,144 DEGs which split into two modules (grey and orange) in a SA-ßGal level-specific manner which enriched for MYC targets and multiple senescence-associated pathways, respectively, consistent with results from freshly purified SA-ßGal-low and -high CD8+ T cells (**see Figure 1**). Interestingly, IL2 treatment induced 1,730 specific DEGs and shared a core of 2,414 DEGs with freshly purified SA-ßGal-low and -high CD8+ T cells, which exhibited 2,289 specific DEGs, indicating a differential response to IL2 stimulation by non-senescent and senescent CD8+ T cells (**Supplementary Figures S6A-C**). Having established this baseline, we perturbed three nodes, KLF5, AP1 (c-JUN), and RUNX2, in the TF networks of both opening and closing chromatin using previously described pharmacological inhibitors SR18662, T5224, and CADD522 ^41–43^, respectively, in SA-ßGal-high CD8+ T cells. We targeted these TFs based on their predicted critical role in the TF networks, as their respective TF families were the most prominently bound top-layer TFs and mediated most interactions with the remaining nodes in the networks (**Figures 5** and **Supplementary Figure S5**). Quantification of the number of binding instances per TF and annotation to their nearest DEGs in both opening and closing chromatin revealed both unique and shared binding instances and DEGs associated to each TF, indicating that these TFs act both individually and as a network in the regulation of senescence-associated gene expression in CD8+ T cells with a magnitude proportional to their number of binding instances (**Figures 6A-B** and **Supplementary Figures S6D-E**). Analysis of bulk transcriptomes of DMSO and inhibitor-treated SA-ßGal-high CD8+ T cells confirmed individual and combinatorial gene regulation by the targeted TFs. For example, genes involved in the unfolded protein and IFNγ responses in module 6 linked to open chromatin were repressed in cells treated with either SR18662 or T5224 but were only modestly affected by CADD522, indicating that these genes are co-regulated primarily by KLF5 and AP1, while genes in module 7, which enriched for hypoxia, complement and apical junction pathways were de-repressed to by treatment with all three inhibitors, suggesting co-regulation of these DEGs by KLF5, AP1 and RUNX2 (**Figures 6C-D**). Similar results were found for DEGs linked to closing chromatin (**Supplementary Figures S6F-G**). In addition, TF inhibitor treatment resulted in both up- and down-regulation of target DEG modules, confirming that TFs act to fine-tune the level of expression of their target genes rather than acting as binary switches (**Figures 6C** and **Supplementary** Figure 6F) ^17,44^. This conclusion is supported by information flow analysis of TF binding and gene expression modules which showed that only a fraction of potential binding sites per TF was involved in the regulation of the perturbed DEG modules (**Figure 6E** and **Supplementary** Figure 6H). Collectively, these results highlight that perturbation of the TF networks is a feasible strategy for the manipulation of the senescence state of CD8+ T cells.

**Figure 6.**
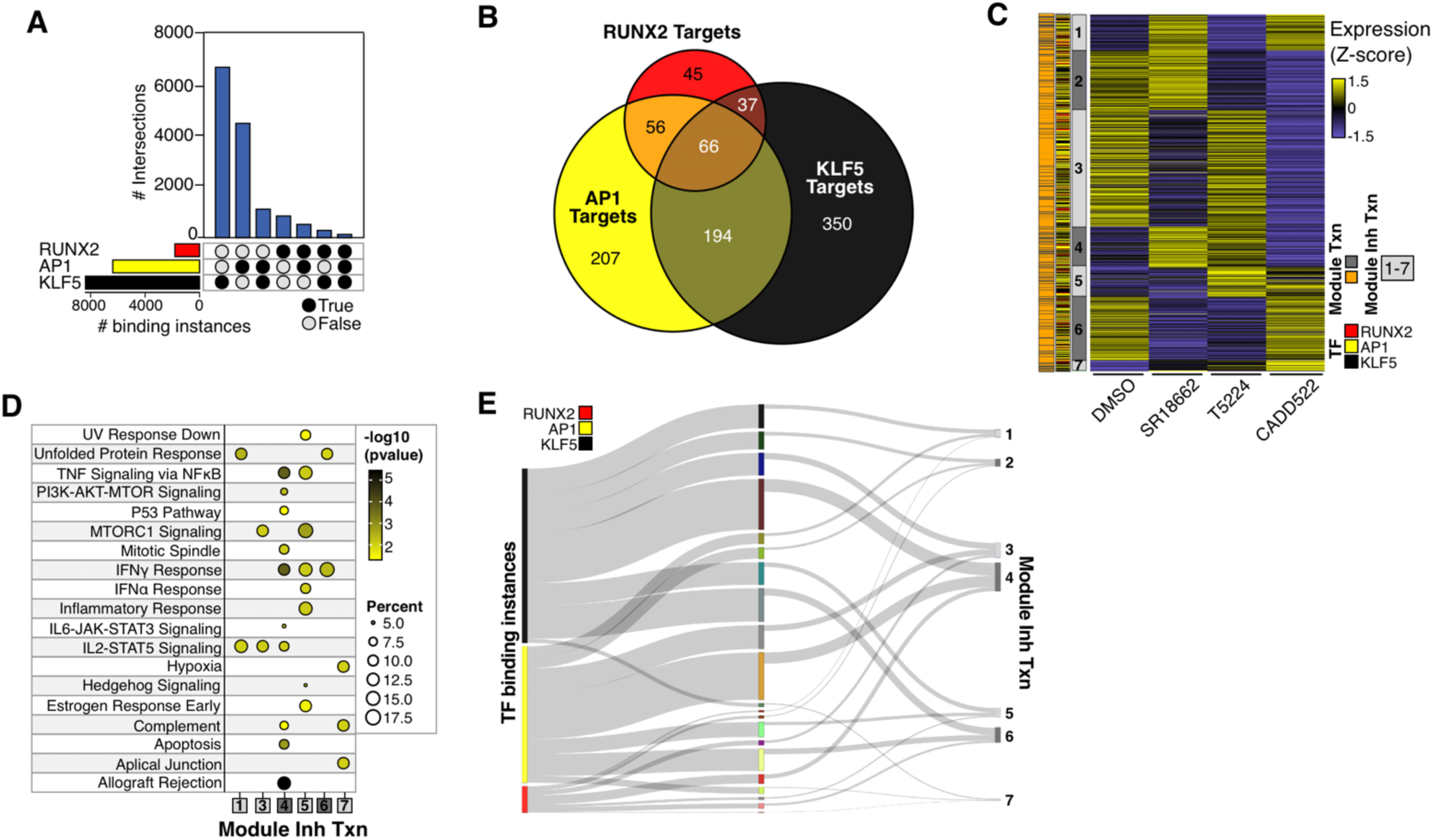
Pharmacological perturbation of the senescence TF network modulates the output of the senescence program of CD8+ T cells. A. Upset plot showing the individual and co-binding instances of KLF5, AP1 and RUNX2 in the senescence-specific open chromatin of CD8+ T cells cultured in the presence of 20U/ml IL2. **B.** Intersections of nearest gene targets for KLF5, AP1 and RUNX2 in the open chromatin of senescent CD8+ T cells **C.** Heatmap showing the effect of KLF5, AP1 and RUNX2 inhibitors (SR18662, T5224, CADD522, respectively) on the expression of the direct, senescence-specific DEGs TF targets linked to open chromatin in SA-ßGal-high CD8+ T cells (row Z-score; n = 956 genes). Treatment was performed in the presence of 20U/ml IL2 and 30 µM of each inhibitor or an equal volume of DMSO as control for 3 days. **D.** Functional overrepresentation analysis map showing significant associations of the MSigDB hallmark gene sets for each expression module shown in C. Circle fill is color coded according to the false discovery rate (FDR)-corrected p value from a hypergeometric distribution test. Circle size is proportional to the percentage of genes in each MSigDB gene set. **E.** River plots showing the proportion of KLF5, AP1 and RUNX2 bound sites linked to the expression of their respective direct gene targets in the open chromatin of senescent CD8+ T cells (represented as the color-coded modules as described in C).

### CD8+ T cell senescence gene signatures as prognostics tools for CAR-T cell therapy and systemic lupus erythematosus

We have previously shown that SA-ßGal-high CD8+ T cells are present across all CD8+ T cell differentiation states and exhibit various features of senescence, including high levels of DNA damage, p16, dysfunctional telomeres, as well as proliferative defects in response to stimulation and enrichment of various dysfunctional CD8+ T cell markers such as PD1, KLRG1 and CD57 ^15^. These results, in conjunction with the extensive epigenomic analyses presented here, indicate that SA-ßGal-high CD8+ T cells have transitioned into a dysfunctional state, which could have a substantial impact on the response to therapeutic treatments as well as disease progression. To test this possibility, we evaluated the effect of senescence on CD8+ T cell expansion under conditions used to generate chimeric antigen receptor (CAR) T cell products by the long-term culturing of bulk CD8+ T cells with low (1.93%-28.8%. n=11 donors) and high (35.6%-81.8%, n=7 donors) senescence (SA-ßGal activity) content. Expectedly, CD8+ T cells with higher initial SA-ßGal activity plateaued at 8.8 +/- 3.9 population doublings by day 30 in culture, whereas CD8+ T cells with low senescence content continued to proliferate until day ∼50, achieving 11.36 +/- 5 population doublings (**Figure 7A**). Subsequently, we generated four CD8+ T cell senescence gene signatures from our transcriptomic datasets (overall, TFs, predicted surface genes and predicted secreted genes) and tested their ability to predict response to CAR-T cell therapy using publicly available, clinically annotated transcriptomic data of naïve CD8+ and CAR-T cell products of a cohort (n=11) of diffuse large B cell lymphomas (DLBCL) ^45^. We found that all four CD8+ T cell senescence gene signatures faithfully identified DLBCLs that were insensitive to CAR-T cell therapy (**Figures 7B-D**; FDR <= 0.07), irrespective of the cell source of the gene expression data (i.e., naïve, CAR-T and a combination of both). Importantly, these results highlight senescence profiling of the starting cellular material used in the generation of CAR-T cell products (naïve and/or memory T cells) as a potentially critical decision-making step during the management of DLBCL. We also tested CD8+ T cell senescence gene signatures as prognostic markers for systemic lupus erythematosus (SLE), an autoimmune disease characterized by autoreactive B cells and aberrant CD8+ T cell function, including organ infiltration and damage ^46^, using published annotated transcriptomic datasets of a cohort of 48 patients ^47^. We found that 3 out of 4 of our CD8+ T cell senescence gene signatures were significantly enriched in the transcriptomes of peripheral CD8+ T cells of patients with active SLE (**Supplementary** Figure 7A) and, to a lesser degree, in patients with inactive SLE (**Supplementary** Figure 7B), indicating a potential role of CD8+ T cell senescence in SLE pathology. In summary, our integrative analysis of CD8+ T cell senescence highlights the potential of senescence-associated gene signatures as prognostic tools in clinical settings with a strong contribution of CD8+ T cells to disease progression and/or response to therapies.

**Figure 7.**
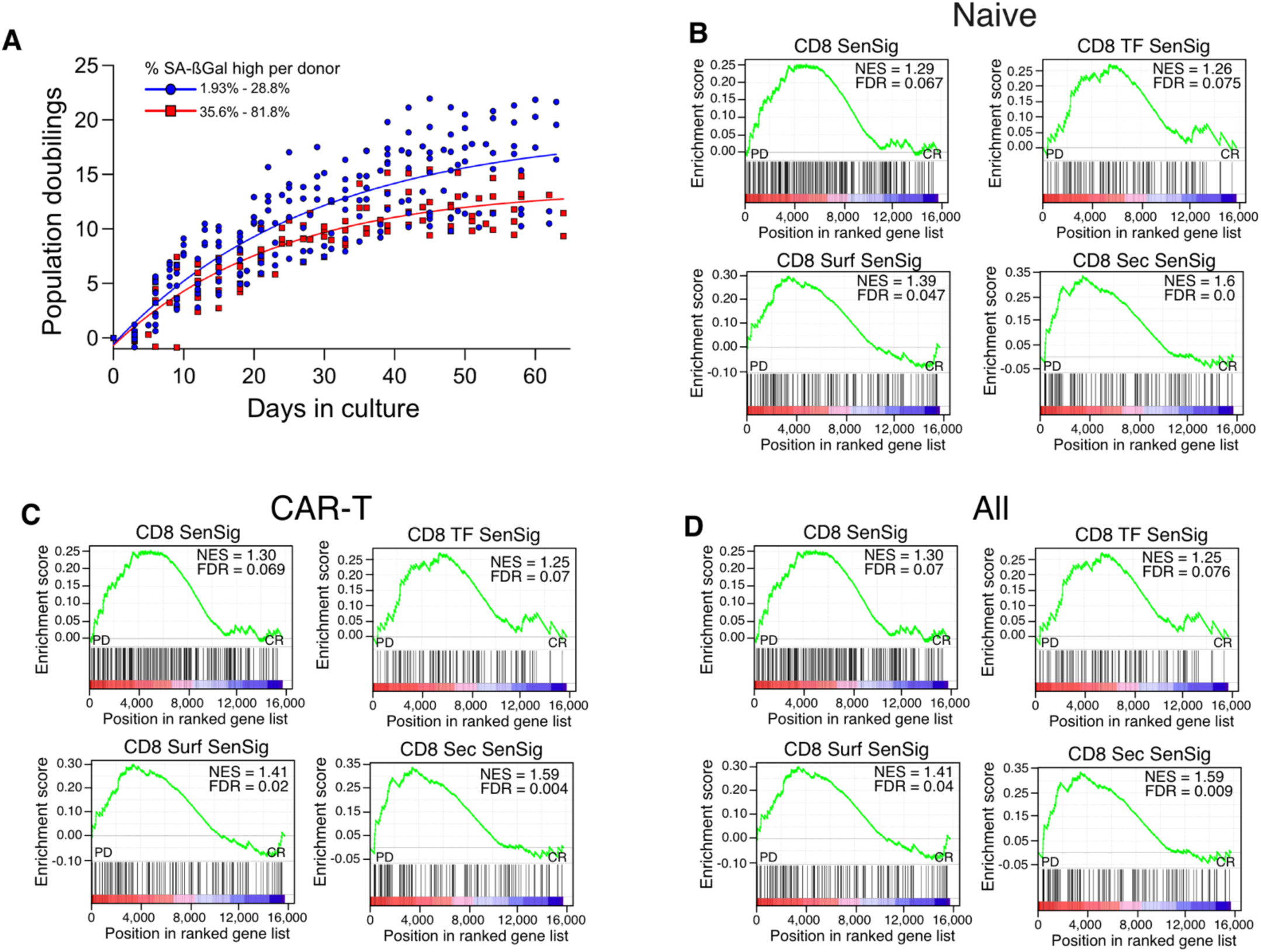
Predictive potential of CD8+ T cell senescence gene signatures of response to CAR-T cell therapy in diffuse large B cell lymphoma (DLBLC). A. Peripheral CD8+ T cells isolated from 19 donors were cultured in Immunocult supplemented with 20U/ml IL2 and stimulated every 15 days using soluble anti CD3/28. CD8+ T cells with 1.93 to 28.8% or 35.6 to 81.8% SA-ßGal high cells were grouped together and labeled as blue or red and cultured into replicative exhaustion. Non-linear fit comparison revealed statistical significance between groups (p < 0.0001). **B-D** Gene set enrichment analyses (GSEA) showing normalized enrichment score (NES) plots and FDR values for overall senescence (SenSig), senescence-specific TF (TF SenSig), surface-predicted (Surf SenSig) and secreted-predicted (Sec SenSig) gene signatures in transcriptomes of naïve CD8+ T cells (B), CAR-T cell products (C) and combined transcriptomes of both (D) of patients with DLBLC (GSE223655) profiled at diagnosis and classified into relapse (n = 9) and no-relapse (n = 11) groups.

## DISCUSSION

The age-associated accumulation of senescent cells in tissues is one of the driving causes of mammalian aging and age-related disease^48,49^. Although senescent cells upregulate the expression of MICA/B ligands and secrete numerous chemokines that facilitate the recruitment of cytotoxic immune cells such as CD8+ T cells for their elimination, senescent cells accumulate systemically with advancing age ^50–52^. These observations suggest that immune cell-mediated elimination of senescent cells becomes impaired with advancing age. Recent evidence highlights a dysfunctional adaptive immune system as a potential cause for the age-associated accumulation in senescent cells. For instance, elimination of the base excision repair enzyme gene *Ercc1* in mouse hematopoietic cells as well as the mitochondrial transcription factor *Tfam* in mouse T cells result in an accelerated aging phenotype, solid organ senescence and dysfunction and multimorbidity ^53,54^. In older humans, both CD4+ and CD8+ T cells acquire features of senescence ^15,55,56^, which is linked to defective immune responses ^57^. However, the gene regulatory mechanisms that promote senescence of CD8+ T cells in aging humans as well as the contribution of senescent CD8+ T cells to disease are poorly understood. Knowledge of the epigenetic mechanisms driving senescence of CD8+ T cells is likely to unveil opportunities for the development of therapies to restore function of damaged CD8+ T cells, potentially increasing healthspan and improving the efficacy of immune cell-based therapies. Using a combination of senescent cell isolation, integrative multiomic profiling and pharmacological perturbations, we defined and validated the TF networks that control the senescence program in CD8+ T cells of a cohort of younger and older donors. Our study lays the foundation for understanding the epigenetic regulation of senescence in CD8+ T cells in otherwise healthy humans and provides a resource for the modulation of the senescence state through targeted perturbation of the TF network.

One key finding of our study is that the acquisition of senescence, as measured by increases in SA-ßGal activity, is the main driver of chromatin accessibility, epigenome and gene expression dynamics of CD8+ T cells, with a minor contribution of chronological age. The transition to the senescence state is a major event in the epigenetic life of CD8+ T cells, as it involved the differential expression of 40% of all detectable TFs and required extensive enhancer remodeling for the implementation of the senescence transcriptional program. These findings are in line with previous work demonstrating the dynamic regulation of the enhancer landscape as a general gene regulatory mechanism required for the establishment of the senescence state in human fibroblasts ^17,58^. However, an important distinction that characterizes CD8+ T cell senescence is the decommissioning of active enhancers as the most prominent chromatin regulatory event, which is heavily associated with the repression of genes characteristic of CD8+ T cell function, including immune surveillance and proliferation. In contrast, widespread enhancer activation is required for the establishment of replicative senescence in cultured human fibroblasts ^17,58^. Thus, although remodeling of the enhancer landscape appears to be a conserved regulatory event during the implementation of the senescence program, defining the chromatin-based mechanisms underlying cell type-specific differences in its execution will be required to manipulate senescence phenotypes in a cell type-specific manner.

Our data also reveal that the senescence competence of CD8+ T cells is precoded at least during early adulthood, as shown by the detection of prebound, chromatin state-specific and hierarchical TF networks of remarkably similar architecture in the CD8+ T cells of both younger and older individuals. The prior association of these TFs with accessible chromatin likely facilitates the rapid activation of the senescence program, independently of the CD8+ T cell differentiation state. Accordingly, we observed near identical activation of the senescence TF network in both age groups, with a few additional TF interactions in the CD8+ T cells of older donors, likely representing more differentiated CD8+ T cells characteristic of this age group. Our results therefore suggest that the ability to undergo senescence is a property of all circulating CD8+ T cells, irrespective of their differentiation state. This mechanism is poised to maximize the regulatory flexibility of potentially long-lived CD8+ T cells ^59^ to undergo senescence when required. In line with this interpretation is our previous quantification of SA-ßGal-positive CD8+ T cells across differentiation states in younger and older donors ^15^, as well as recent evidence characterizing TF activity across CD8+ T cell differentiation states in aging humans ^21,23^. Our comprehensive analysis of epigenomic data confirmed the hierarchical architecture of the CD8+ T cell senescence TF networks, consistent with our previous work on senescent fibroblasts ^16,32^, and provides a logical platform for the manipulation of senescence in CD8+ T cells. Accordingly, using a pharmacological inhibition approach and leveraging the conservation of the senescence TF network in CD8+ T cells of both younger and older humans, we precisely modulated the transcriptome of senescent CD8+ T cells by targeting key nodes c-JUN (AP1), RUNX2 and KLF5. Collectively, these findings provide proof of principle that targeting the senescence TF network is a viable approach to diminish the potentially damaging effects of senescent CD8+ T cells.

Another important implication of our study is the potential contribution of senescent CD8+ T cells to the effectiveness of CAR-T cell therapy and autoimmune disease. We observed robust enrichment of CD8+ T cell senescence gene signatures derived from our datasets in the starting cellular material (naïve CD8+ T cells) and the CAR-T cell product of a cohort of patients with DLBLC who were insensitive to CAR-T cell therapy. This, in combination with our data showing the decreased proliferative potential of bulk CD8+ T cells with a higher senescence burden under conditions of CAR-T cell product expansion, highlights senescence as a critical block to the development of effective CAR-T cell products. Similarly, our CD8+ T cell senescence gene signatures were highly enriched in the peripheral blood of individuals with active and inactive SLE. Although the role of senescent CD8+ T cells in SLE pathogenesis and progression is presently unknown, their highly inflammatory nature, along with the reduced expression of functional genes, including those involved in immune surveillance, is consistent with the presence of previously described overreactive inflammatory CD8+ T cell subpopulations in patients with active SLE ^60^. Future work will be required to establish the molecular mechanisms leading to CD8+ T cell senescence in the context of autoimmunity and CAR-T cell therapy effectiveness.

In conclusion, our study provides a comprehensive foundation of the gene regulatory mechanisms governing senescence in CD8+ T cells and provides a logical platform for its modulation through rational targeting of the senescence TF network. Our study also highlights senescent cell isolation followed by multiomic profiling as a powerful strategy to study the contribution of immune cell senescence to aging and age-related disease.

## METHODS

### CD8+ T cell enrichment

The Institutional Review Board of New Jersey Medical School approved this study (ID Pro2020000006, approved May 14, 2024). Peripheral blood mononuclear cells (PBMCs) were obtained from the blood of consenting healthy donors (aged 20–60 years) collected in heparinized tubes (BD Biosciences, Franklin Lakes, NJ). PBMCs were isolated using density gradient centrifugation with lymphocyte separation medium (Corning, Manassas, VA). Briefly, heparinized blood was diluted 1:1 with Hanks’ Balanced Salt Solution (HBSS), (Corning, Manassas, VA), layered over lymphocyte separation medium at a 2:1 ratio, and centrifuged at 400g for 30 minutes with reduced deceleration (deceleration factor of 2). The buffy coat containing PBMCs was carefully harvested and processed further. CD8+ T cells were enriched from PBMCs using negative selection with the EasySep™ Human CD8+ T Cell Isolation Kit (StemCell Technologies, Cambridge, MA), following the manufacturer’s protocol. The purity of the isolated CD8+ T cells was assessed via flow cytometry by staining for surface markers PerCP/Cyanine7-CD3 (HIT3a), and PE-CD8 (RPA-T8), BioLegend, San Diego, CA. Briefly, all surface staining was conducted at 4C for 20 minutes in 1xPBS+ 2% FCS whilst protected from light. Samples were subsequently washed 1xPBS+ 2% FCS and kept on ice before being acquired on the BD LSRFortessa X-20 (BD Biosciences, Franklin Lakes, NJ). Samples consistently achieved >90% CD3+CD8+ double-positive cells.

### Surface, SA-βGal staining and FACS

Isolated CD8+ T cells were resuspended in RPMI-1640 medium containing 10% fetal bovine serum (FBS, Atlanta Biologicals) and 1× penicillin/streptomycin (Corning, Manassas, VA). The Cellular Senescence Detection Kit-SPiDER-βGal (Dojindo Molecular Technologies, Inc, Rockville, MD) was used to identify senescent CD8+ T cells as previously described. Briefly, CD8+ T cells were incubated with a 1:1000 dilution of bafilomycin A-1 for 30 min before the addition of 1:1000 dilution SPiDER-βGal for an additional 30 min all whilst at 37C, 5% CO2. SA-βGal stained cells were subsequently washed with 1x PBS (Corning, Manassas, VA) and resuspended in 1xPBS + 2% FBS. For ATAC and RNA sequencing, enriched CD8+ T cells were sorted based on live, 4′,6-diamidino-2-phenylindole (DAPI (BioLegend, San Diego, CA) DAPI-, SA-βGal intensity. For CUT&TAG experiments, CD8+ T cells were sorted based on the SA-βGal intensities of DAPI-, CD8+ T cells. All cells were sorted using the BD FACSAria Fusion (BD Biosciences, Franklin Lakes, NJ).

### In vitro expansion

Purified CD8+ T cells (>90% CD3+CD8+) were cultured at 37°C in a humidified atmosphere with 5% CO₂. Cells were maintained in ImmunoCult™ Expansion Medium (StemCell Technologies, Cambridge, MA) supplemented with 20 U/mL IL-2 (StemCell Technologies). T cells were stimulated every 15 days using soluble anti-CD3 and anti-CD28 antibodies (StemCell Technologies). The growth medium was replenished every 2–3 days to ensure optimal cell viability and proliferation. Cell growth was monitored by counting viable cells using a hemocytometer. Population doublings (PD) were calculated using the formula PD=3.32∗[log(N final)-log(N initial)], where N final is the number of cells recovered and N initial denotes the number of cells initially seeded at each flask.

### RNA-seq

RNA from sorted CD8+ fSA-ßGal low and high from young and old donors (7 each) was purified using a Macherey-Nagel RNA XS Plus kit according to the manufacturer’s instructions (Macherey-Nagel, Duren, Germany). RNA integrity was evaluated in a Bioanalyzer 2100 system, and only RNA with an integrity of number of >= 7 was used for library preparation. Libraries were constructed using the SMARTer Stranded V2 (TakaraBio) according to the manufacturer’s instructions (TakaraBio, Mountain View, CA). Paired-end sequencing was performed on an Illumina HiSeq 2500 instrument. At least 40 million reads per sample (20 million per strand) were obtained and used for downstream analyses.

### ATAC-seq

The transposition reaction and library construction were performed as previously described in ^26^. Briefly, 50,000 cells from sorted CD8+ fSA-ßGal low and high from young and old donors (4 each) were collected, washed in 1× in PBS, and centrifuged at 500 × g at 4 °C for 5 min. Nuclei were extracted by incubating cells in nuclear extraction buffer (containing 10 mM Tris-HCl, pH 7.4, 10 mM NaCl, 3 mM MgCl2, 0.1% IGEPAL CA-630) and immediately centrifuged at 500 × g at 4 °C for 5 min. The supernatant was carefully removed by pipetting, and the transposition was performed by resuspending nuclei in 50 μl of Transposition Mix containing 1× TD Buffer (Illumina, San Diego, CA) and 2.5 μl Tn5 (Illumina) for 30 min at 37 °C. DNA was extracted using a Qiagen MinElute kit (Qiagen, Germantown, MD). Libraries were produced by PCR amplification (12 cycles) of tagmented DNA using an NEB Next High-Fidelity 2× PCR Master Mix (New England Biolabs, Ipswich, MA). Library quality was assessed using an Agilent Bioanalyzer 2100 (Applied Biosystems, Santa Clara, CA). Paired-end sequencing was performed in an Illumina Hiseq 2500 system. Typically, 30–50 million reads per library were required for downstream analyses.

### CUT&Tag

We performed CUT&Tag on 75,000–100,000 sorted CD8+ fSA-ßGal low and high from young donors (n=3) with the CUTANA CUT&Tag kit (Epicypher) using the following antibodies: H3K4me1 (Epicypher: 13–0057), H3K27me3 (Epicypher: 13–005), H3K27ac (Active Motif: 39,133) and rabbit IgG as a negative control (Epicypher: 13–0042), according to the manufacturer’s instructions. Libraries were amplified for 16 cycles and quality assessed using a TapeStation 4200 instrument. Libraries were paired-end sequenced on a Novaseq-X instrument. 10–20 million reads per library were used for downstream analyses.

### Preprocessing of high-throughput sequencing data

Paired-end (RNA-seq, CUT&Tag and ATAC-seq) reads were processed as we previously described ^32^ aligned to the GRCh38.d1.v1 version of the human genome using bowtie2 ^61^ using the local mode (RNA-seq) or with the following parameters for ATAC-seq and CUT&Tag: -*N 0 --no-mixed --no-discordant --maxins 2000 -x --trim3 8*. Low-quality reads and adapters were removed using fastq-mcf v.1.0.5 and cutadapt. Alignments were further processed using samtools v.1.1.1, and PCR and optical duplicates were removed with PicardTools v.2.2.2. Enriched regions ATAC-seq were identified using MACS v.2.2.7.1 ^62^ using a relaxed q-value threshold of 0.01. Master peaksets were constructed using a custom bedops script that merges common peaks between samples as we previously described ^16^. CUT&Tag data were processed using chromstaR (see below). For RNA-seq samples, reads were counted using summarized overlaps and normalized with DESeq2 for visualization.

### Self-organizing maps (SOMs)

SOM expression portraits were generated using the unsupervised machine learning method deployed in oposSOM ^25^. Metagenes were visualized in a 60 x 60 grid of rectangular topology, wherein expression portraits are projected by metagene distance matrix similarity using a logarithmic fold-change scale. To verify that expressions portraits from GM21 fibroblasts escaping OIS differ from time-matched controls, we implemented the D-clustering feature of oposSOM, which reveals clusters of differentially expressed genes based on SOM units with local maxima relative to the mean Euclidean distance to their neighbors.

### Differential expression and accessibility

Differentially accessible regions (DARs) and differentially expressed genes (DEGs) throughout the time course were identified from ATAC-seq and RNA-seq data, respectively using DESeq2. Raw reads are internally normalized by DESeq2 using the median of ratios method ^63^. Reads per peak (ATAC-seq, using the master peakset as feature input) or exon (RNA-seq, using the GRCh38.107 genome model) were quantified using the *summarizeOverlaps* package and peaks/genes with at least 10 reads in at least 10 libraries were kept. Correction of batch effects was performed with limma ^64^ using the nucleic purification date (batch) as the surrogate variable. Data transformation (regularized-log [rld] transformation), exploratory visualization (PCA and hierarchical clustering) and differential expression analysis were performed with DESeq2 using the default parameters as previously described ^65^ and base R functions. We focused on highly significant DARs and DEGs by using an adjusted *p*-value filter of 0.05.

### WGCNA and integration of chromatin accessibility with differential expression

Differentially expressed genes (DEGs) and accessible regions (DARs) identified with DESeq2 were used as input for unsupervised clustering using WGCNA. We used the “signed” option with default parameters, except for the soft thresholding power which was set to 17 for RNA-seq data and 6 for ATAC-seq data. The minimum size for the DEG and DAR modules was set to 200 features for the initial set of modules, which were then merged by a dissimilarity threshold of 0.5 for RNA-seq data and 0.2 for ATAC-seq data. For RNA-seq, WGCNA modules were functionally profiled using clusterProfiler ^66^ using the Molecular Signatures Database Hallmark gene sets ^67^. Statistical significance was calculated by a hypergeometric test with a cut-off of an adjusted *p*-value of <= 0.1 with Benjamini–Hochberg correction. Integration of DARs with DEGs was achieved by annotation of DARs to the nearest gene using ChIPSeeker ^68^ followed by filtering against the list of DEGs.

### Chromatin state transitions

We analyzed the genome-wide combinations of H3K4me1, H3K27ac, H3K27me3 and ATAC-seq signals modeling senescence as a transition from fSA-ßGal low to fSA-ßGalusing the chromstaR package essentially as we previously described ^32^. The algorithm was run on differential mode and configured to partition the genome in 200 bp non-overlapping bins and count the number of reads of histone modification/ATAC-seq mapping into each bin and modeled using a univariate HMM based on a two-component mixture, the zero-inflated binomial distribution. Subsequently, a multivariate HMM assigns every bin in the genome to one of the multivariate components considering 2(2 cell states x 4 genomic enrichment variables [histone modifications and ATAC-seq]) possible states. We focused on robust transitions of an enrichment score of >= 1.5 for further analysis. Integration with gene expression data was achieved by annotating the nearest DEGs with ChIPSeeker ^68^ to the 10 most frequent chromatin state transitions and monitoring their expression using violin plots. The association between chromatin state transitions and DEG modules was performed through Correspondence analysis (CA) and visualized on an asymmetric biplot after filtering for the top contributing chromatin state transitions (square cosine >0.5).

### Heatmap and metaprofile visualizations of ATAC-seq and CUT&Tag datasets

ATAC-seq and CUT&Tag alignments were normalized using deeptools v3.3.1100103 ^69^ using the RPGC approach to obtain 1X coverage (bamCoverage -b–normalizeUsing RPGC–effectiveGenomeSize 2864785220 –ignoreDuplicates–binSize 10 –verbose -o).

### Transcription factor footprinting

Footprinting, TF motif enrichment, and differential binding activity were performed as we previously described ^32^ with HINT-ATAC ^70^ using the JASPAR position weight matrix database for vertebrate TFs ^71^ on merged ATAC-seq datasets, focusing on differentially accessible chromatin in fSA-ßGal CD8+ T cells. TF co-binding was assessed across <=500bp bins of differentially accessible chromatin using a hypergeometric test for each pairwise co-binding possibility for each condition (fSA-ßGal low and high) and adjusted *p*-values for multiple testing using Bonferroni correction.

### Transcription factor networks

We built TF hierarchical networks as previously described in ^17^, at opening and closing DARs. Each identified TF is represented as a node, connected by directed edges representing co-binding events along the chromatin and during the transition from fSA-ßGal low to fSA-ßGal high. An edge with TF A as the source and TF B as the target indicates that TF A binds to the same chromatin regions as TF B at the same or at a previous time point. For each TF, we computed their normalized total number of bound regions and dynamicity and represented those properties in the networks as node size and node color, respectively. Each edge is associated with a weight in the interval [0, 1], representing the fraction of TF B binding instances previously occupied by TF A. We filtered the networks for edges with a weight higher than 0.15 and performed a transitive reduction to simplify the obtained networks while keeping essential topological features. Nodes included in the same strongly connected component (SCC), i.e., connected both by incoming and outgoing paths, were merged into a single node. We represented the identified transcriptomic module distribution for TFs included in the same SCC as Doughnut Charts.

### Pharmacological perturbation of transcription factor networks

Sorted SA-βGal high CD8+ T cells were young donors (n=3) cultured for three days in RPMI-1640 medium supplemented with 10% fetal bovine serum (FBS; Atlanta Biologicals), 1× penicillin/streptomycin (Corning), and 20 U/mL IL-2. Cells were treated with either a vehicle control (DMSO) or one of the following inhibitors at a concentration of 30 µM: SRI8662, T-5224, or CADD522 (Selleckchem, Houston, TX), targeting KLF5, AP1, and RUNX2, respectively. Following the three-day treatment period, cells were collected and RNA and nuclei isolated for RNA-seq and ATAC-seq as described above. To evaluate the effect of pharmacological inhibition of TF network nodes, we annotated KLF5, AP1 and RUNX2 footprints in DARs of fSA-ßGal high CD8+ T cells to the nearest DEG as defined above to identify their potential gene targets, and the effect of TF inhibition on target DEG expression in both opening and closing chromatin relative to DMSO-treated cells was evaluated by clustering the expression levels of TF target DEGs using WGCNA as above (soft-threshold power of 6 and merge distance of 0.4 for opening chromatin; soft-threshold power of 12 and merge distance of 0.4 for closing chromatin) and visualized on a heatmap.

### GSEA

GSEA of 4 CD8+ T cell senescence gene signatures derived from our transcriptomic datasets was performed on transcriptomes (GEO: GSE223655) of naïve CD8+ T cells and CAR-T cell products from diffuse large B cell lymphoma (DLBCL) patients undergoing immunotherapy were phenotypically classified as ‘progressive disease or ‘complete response’ and transcriptomes from peripheral CD8+ T cells from control and individuals with active and inactive systemic lupus erythematosus (SLE) (GEO: GSE97264) using the GSEA GUI version 4.2.2. Probe sets were collapsed to the gene level using the correlation-based approach. The correlation of probe sets representing the same gene was computed to decide whether to average probe sets (c > 0.2) or to use the probe set with the highest average expression across samples (c ≤ 0.2). Probe sets without known annotation were removed. The signal-to-noise ratio (μA – μB)/(σA + σB) (μ represents the mean, σ the standard deviation) was used as a ranking metric and statistics based on 500 gene set permutations.

## ACKNOWLEDGMENTS

This research was supported by start-up funds to R.I.M-Z and a grant from the National Institute on Aging of the National Institutes of Health (R21AG067368) to UH and PFB. The content is solely the responsibility of the authors and does not necessarily represent the official views of the National Institutes of Health.

---

**Figure S1.**
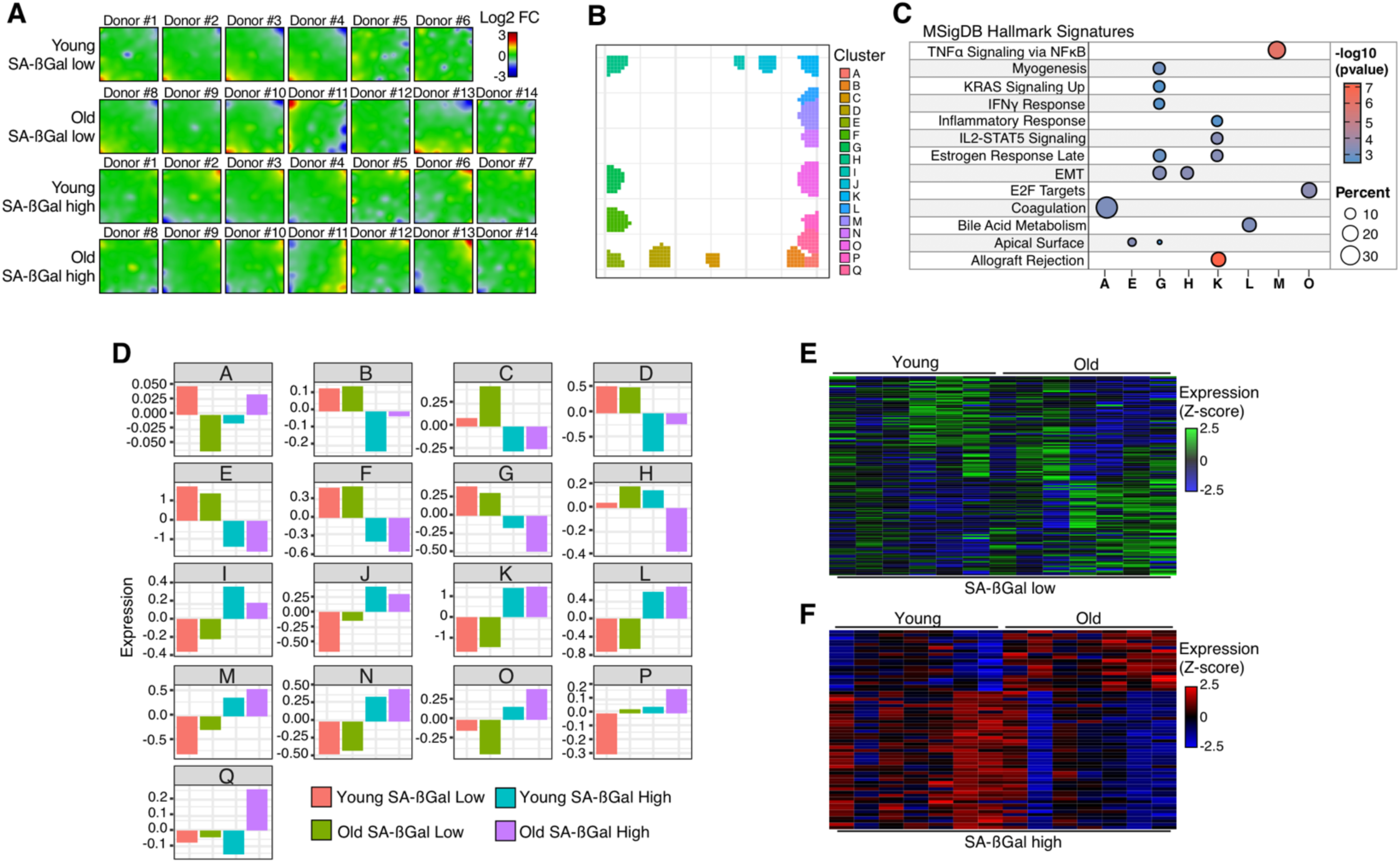
Transcriptional landscape of senescent CD8+ T cells. A. Individual SA-ßGal-low and SA-ßGal-high CD8+ gene expression SOMs. Individual younger and older donors are labelled. **B.** D-cluster projection showing SOM spots with significant changes in metagene expression. **C.** Functional overrepresentation analysis map showing significant associations of the MSigDB hallmark gene sets for the indicated spots from the D-cluster projection in B. Circle fill is color coded according to the false discovery rate (FDR)-corrected p value from a hypergeometric distribution test. Circle size is proportional to the percentage of genes in each MSigDB gene set. **D.** Averaged D-cluster metagene expression changes (histograms) identified in C. Histograms are color coded according to the donor age and SA-ßGal level. **E, F.** Heatmap showing individual donor profiles of age specific DEGs in SA-ßGal-low (E) and SA-ßGal-high (F) CD8+ T cells. A-F were generated from transcriptomic data from SA-ßGal-low and SA-ßGal-high CD8+ T cells of seven younger and seven older donors (A-F).

**Figure S2.**
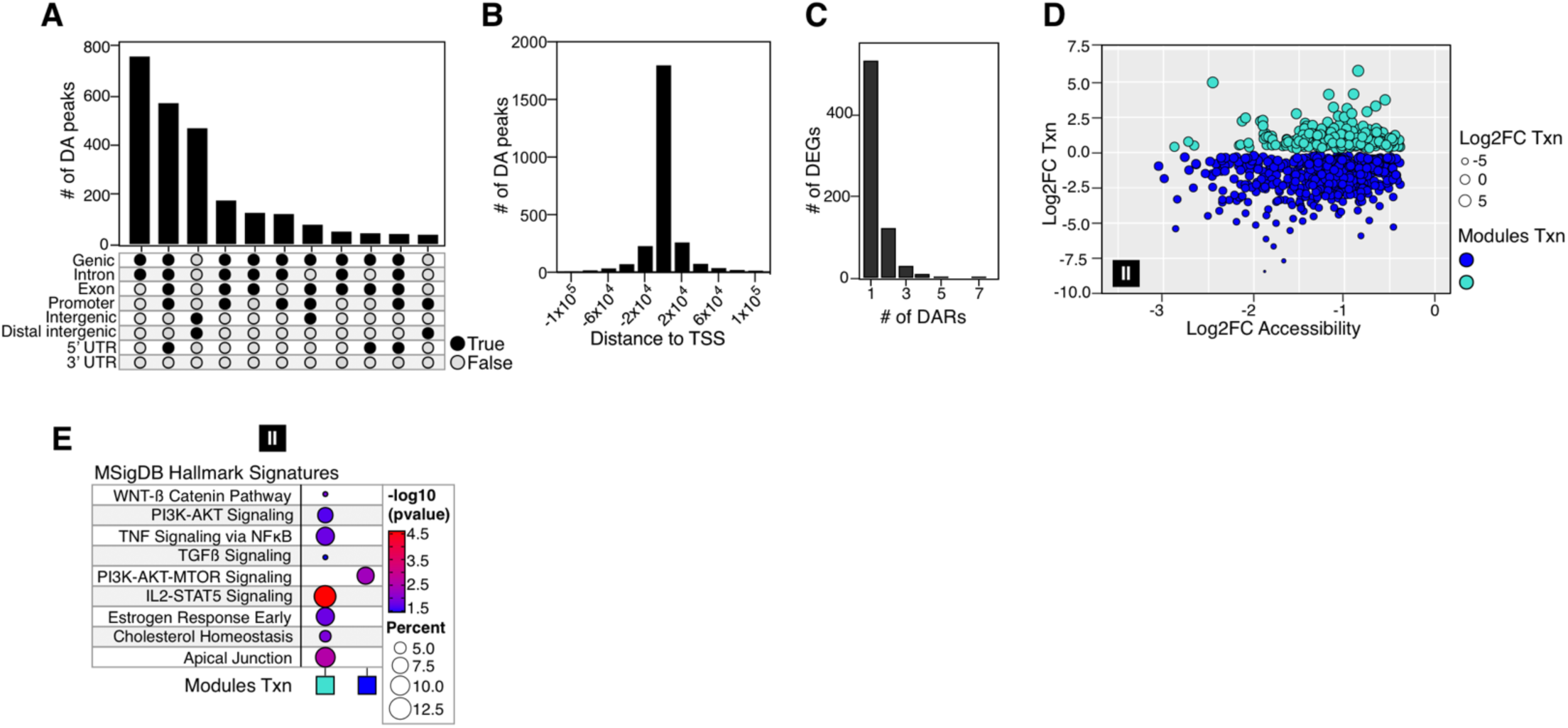
Senescence and not age drives changes in chromatin accessibility of CD8+ T cells. A. UpSet plot showing the frequency distribution of closing DA peaks (black module) with the indicated genomic features. **B.** Frequency distribution showing the occurrence of closing DA peaks relative to the TSS of the nearest annotated gene. **C.** Histogram plot showing the number of DEGs linked to closing DA peaks. **D.** Correlation between chromatin accessibility and expression of DEGs linked to closing DA peaks. Individual DEGs from the blue and turquoise modules from 1C are shown as color coded circles. Circle size is proportional to the log2 fold change in expression. **E.** Functional overrepresentation analysis map showing significant associations of the MSigDB hallmark gene sets for the indicated DEGs linked to opening DA peaks. Circle fill is color coded according to the false discovery rate (FDR)-corrected p value from a hypergeometric distribution test. Circle size is proportional to the percentage of genes in each MSigDB gene set. Chromatin accessibility data (A-D) were generated from SA-ßGal-low and SA-ßGal-high CD8+ T cells from four younger donors and four older donors and integrated with the transcriptomic data from SA-ßGal-low and SA-ßGal-high CD8+ T cells of seven younger and seven older donors (C-E).

**Figure S3.**
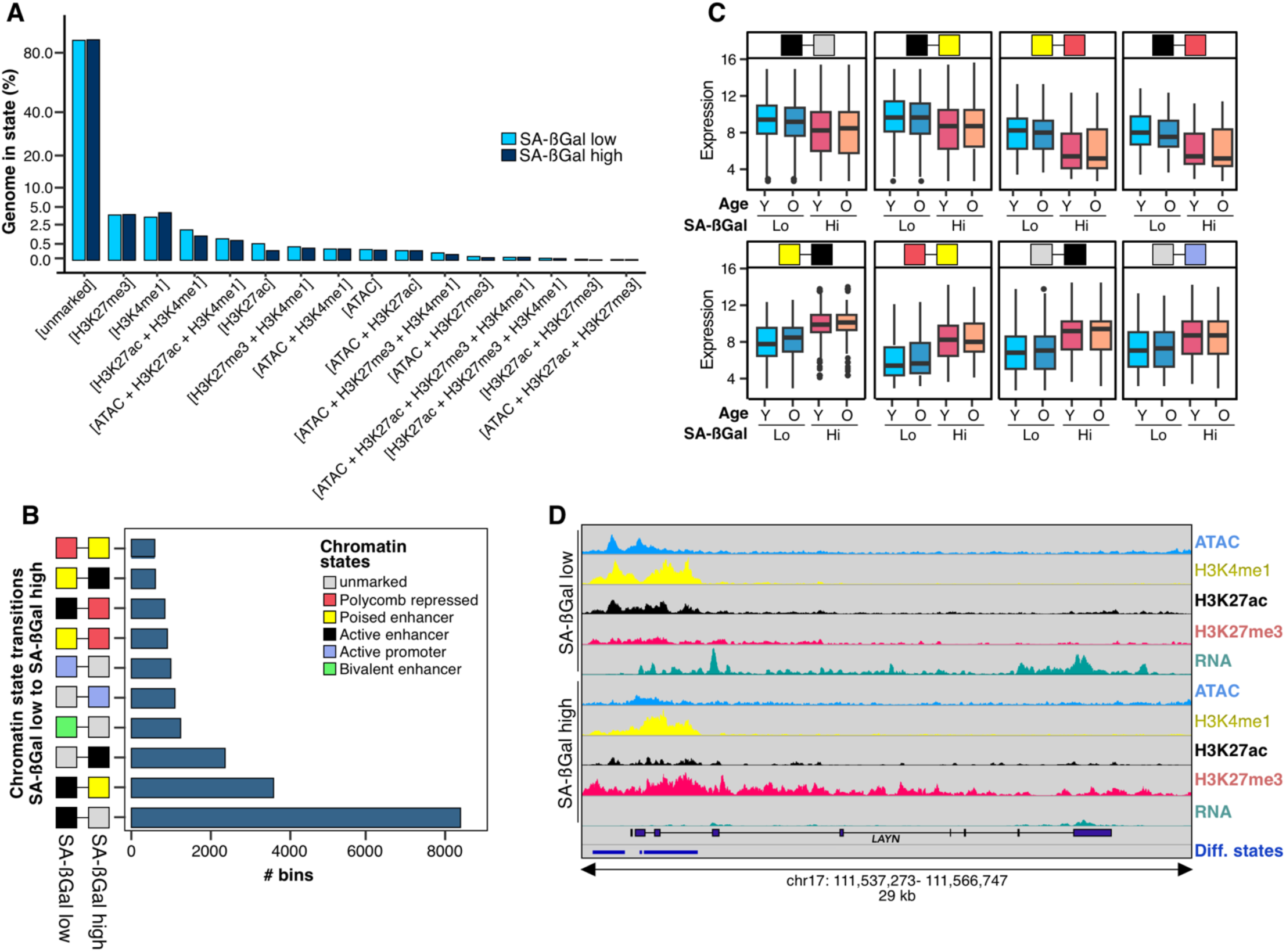
Enhancer remodeling drives the transition to senescence in CD8+ T cells. A. Histograms depicting the percentage of the mapped genome enriched for the indicated combinations H3K4me1, H3K27ac, H3K27me3 Cut&Tag and ATAC-seq signals in SA-ßGal-low and SA-ßGal-high CD8+ T cells. **B.** Quantification of the 10 most common chromatin state transitions as CD8+ T cells transition into senescence. **C.** Integration of eight representative chromatin state transitions with nearby gene expression (row Z-score boxplots). . Thick line in boxplots indicates the median. Edges correspond to the first and third quartiles. Whiskers extend from the edges to 1.5 X the interquartile range to the highest and lowest values. **D.** Representative genome browser snapshot of the *LAYN* locus signal tracks for ATAC-seq, H3K4me1, H3K27ac and H3K27me3 in SA-ßGal-low and SA-ßGal-high CD8+ T cells. Cut&Tag data (A-D) were generated from SA-ßGal-low and SA-ßGal-high CD8+ T cells from three younger donors and integrated with the transcriptomic data from SA-ßGal-low and SA-ßGal-high CD8+ T cells of seven younger and seven older donors (C, D).

**Figure S4.**
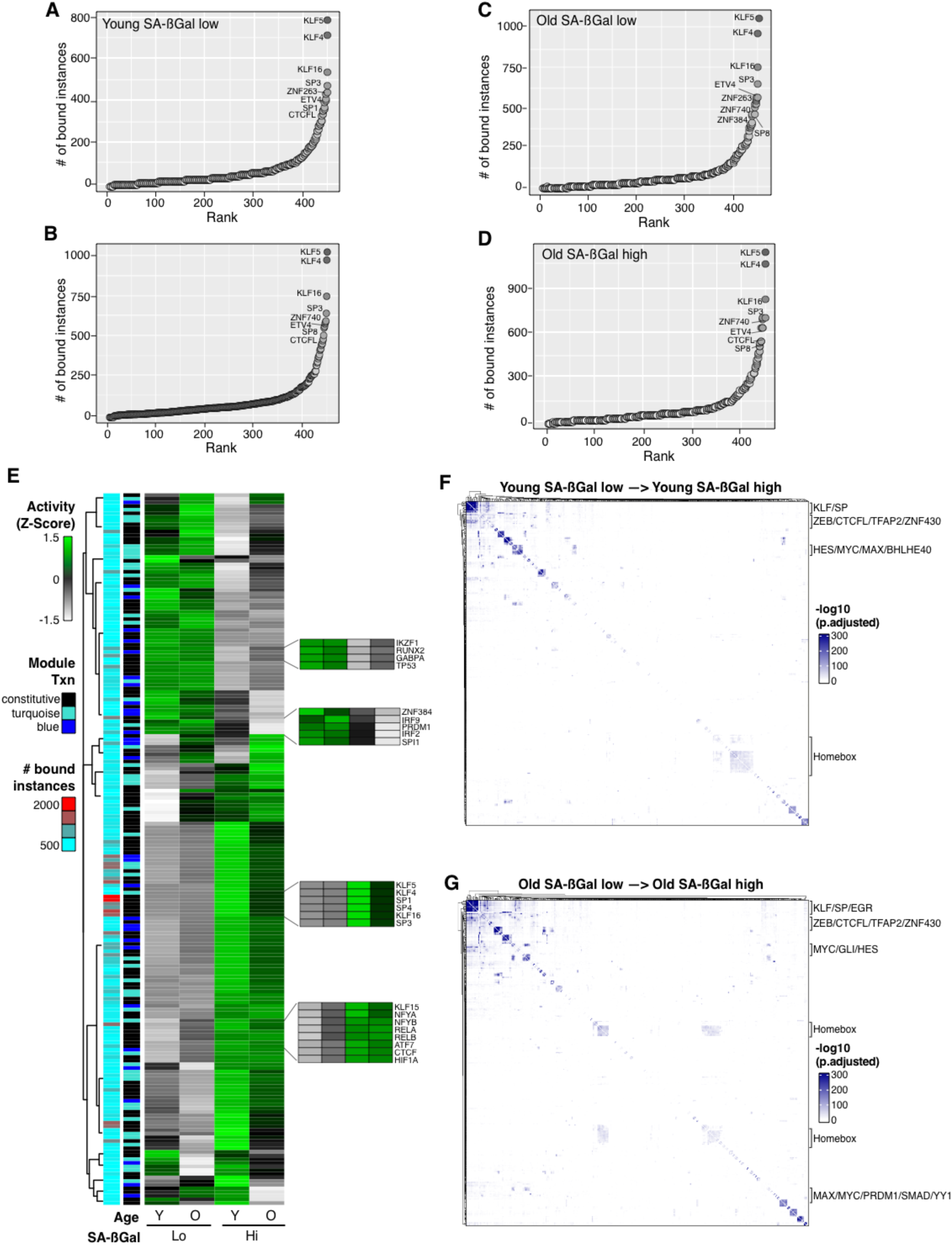
Transcription factor network dynamics of senescent CD8+ T cells. A, B, C, D. Rank plots showing the summed binding instances per TF in the senescence-specific closed chromatin of SA-ßGal-low (A, C) and SA-ßGal-high (B, D) of CD8+ T cells of younger and older donors. The TFs with the most binding instances are labelled. **E.** Heatmap of differential TF chromatin binding activity (row Z-score) at senescence-specific closing chromatin of SA-ßGal-low and SA-ßGal-high CD8+ T cells from younger and older donors. The left annotation heatmaps show the number of bound instances per TF (cyan to red) and their gene expression (TXN) category (i.e., constitutively [black] or differentially expressed according to the module color shown in Figures 1D and E. Insets: chromatin binding activity of age- and cell state-specific TFs. **F, G.** TF co-binding interaction matrices in the senescence-specific closed chromatin of CD8+ T cells from younger (F) and older (G) donors transitioning into senescence. All binding instances across the two cell states (non-senescent and senescent) are collapsed onto the matrices and clustered using Ward’s aggregation criterion. The corresponding q values of the interactions are projected onto the clustering and represented in a color scale defined by their significance using a hypergeometric distribution test. A-G were generated from pooled ATAC-seq datasets from SA-ßGal-low and SA-ßGal-high CD8+ T cells from four younger and four older donors.

**Figure S5.**
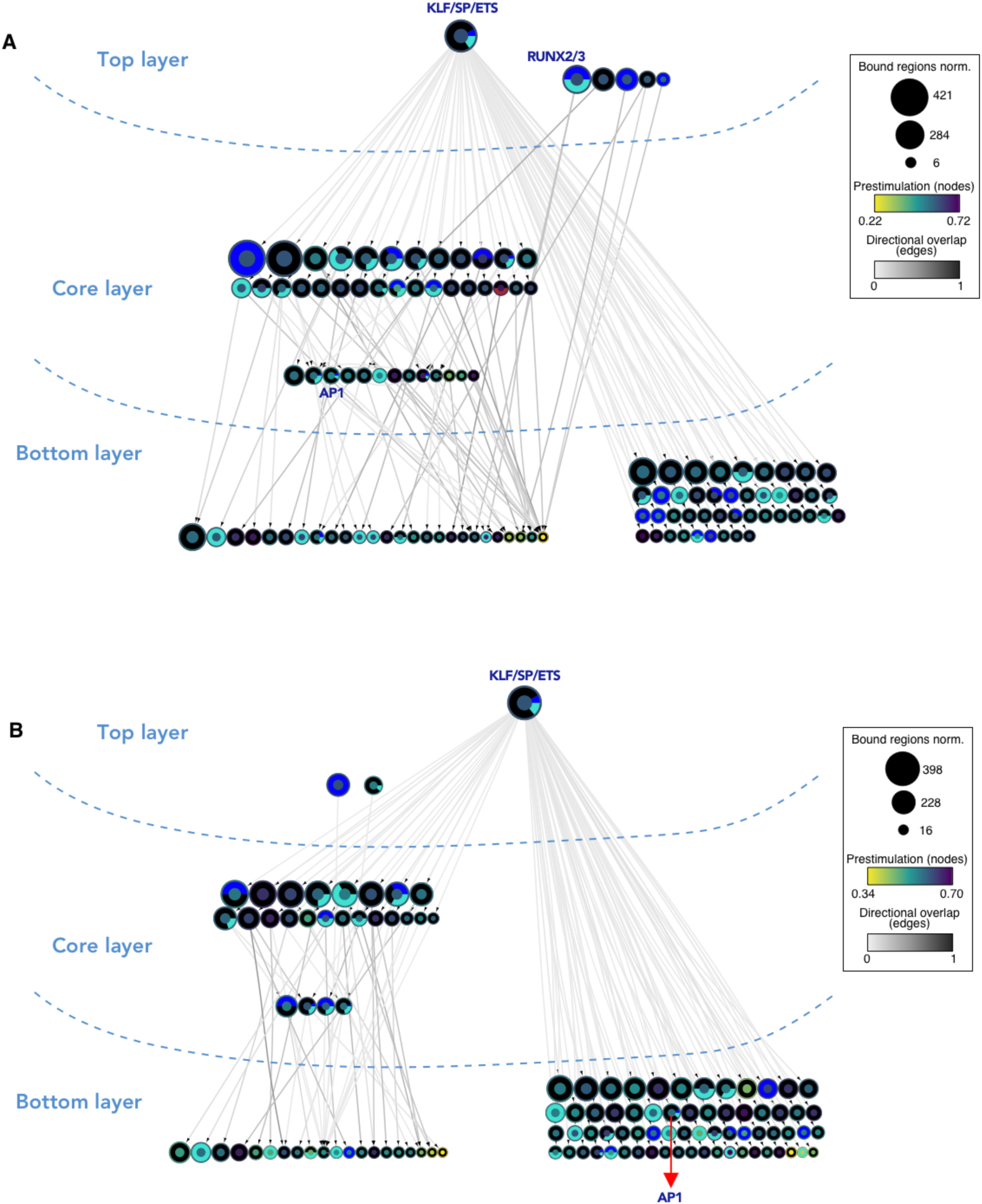
Distinct hierarchical TF networks control the senescence program of CD8+ T cells. A, B. TF networks for the senescence-specific closed chromatin module (Figure 2B, black module) in CD8+ T cells from younger (A) and older (B) donors transitioning into senescence. TFs (nodes) are represented as circles. Oriented edges (arrows) connecting nodes indicate that at least 15% of the regions bound by a given TF in the bottom and core layers were bound by the interacting TF in the core and top layers, respectively, in SA-ßGal-low CD8+ T cells. Nodes represent strongly connected components (SCCs) (i.e., regulons of multiple TF co-binding interactions) to facilitate visualization. The fill color of the node’s inner circle is based on the normalized dynamicity (prestimulation) of TFs. The fill color of the outer ring indicates whether the TF is constitutively expressed (black) or belongs to a transcriptomic module as described in Figure 1D. The node’s size is proportional to the bound regions by a given TF(s). Each network has three layers: (1) the top layer with no incoming edges, (2) the core layer with incoming and outgoing edges, and (3) the bottom layer with no outgoing edges. Bold labels indicate TFs targeted with pharmacological inhibitors (see Figures 6 and S6). Networks were generated from pooled ATAC-seq datasets from SA-ßGal-low and SA-ßGal-high CD8+ T cells from four younger and four older donors.

**Figure S6.**
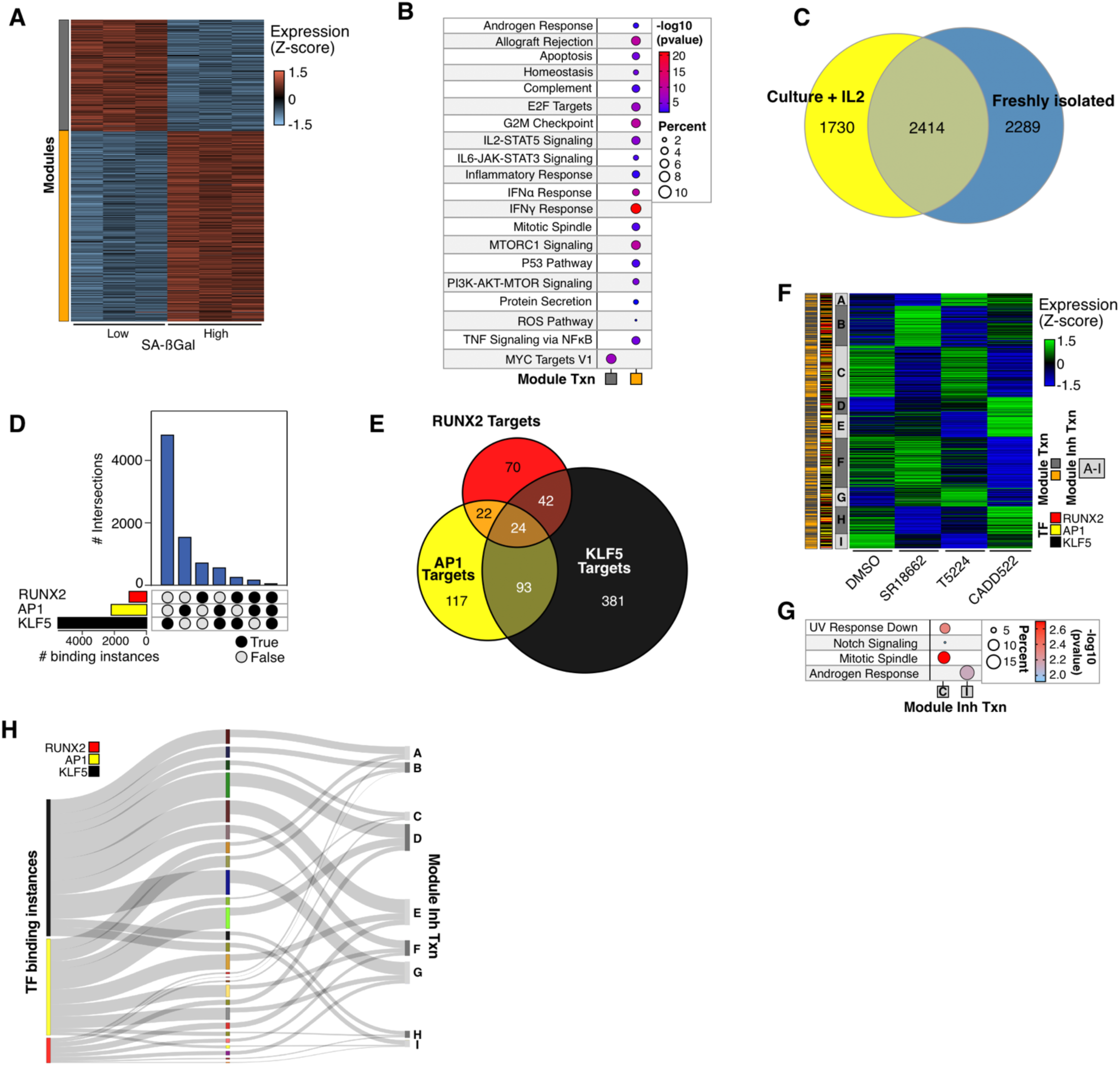
Pharmacological perturbation of the senescence TF network modulates the output of the senescence program of CD8+ T cells. A. Individual heatmaps of color-coded modules of differentially expressed genes (DEGs) in SA-ßGal-low and SA-ßGal-high CD8+ T cells from younger donors stimulated with 20U/ml IL2. **B.** Functional overrepresentation analysis map showing significant associations of the MSigDB hallmark gene sets for each expression module shown in A. Circle fill is color coded according to the false discovery rate (FDR)-corrected p value from a hypergeometric distribution test. Circle size is proportional to the percentage of genes in each MSigDB gene set (B,G). **C.** Euler diagrams showing the DEG overlaps between SA-ßGal-low and SA-ßGal-high freshly isolated CD8+ T cells and stimulated with IL2 for 3 days. **D.** Upset plot showing the individual and co-binding instances of KLF5, AP1 and RUNX2 in the senescence-specific closed chromatin of CD8+ T cells cultured in the presence of 20U/ml IL2. **E.** Intersections of nearest gene targets for KLF5, AP1 and RUNX2 in the closed chromatin of senescent CD8+ T cells **F.** Heatmap showing the effect of KLF5, AP1 and RUNX2 inhibitors (SR18662, T5224, CADD522, respectively) on the expression of a subset of senescence-specific DEGs linked to closed chromatin in SA-ßGal-high CD8+ T cells (row Z-score; n = 749 genes). Treatment was performed in the presence of 20U/ml IL2 and 30 µM of each inhibitor or an equal volume of DMSO as control for 3 days. **G.** Functional overrepresentation analysis map showing significant associations of the MSigDB hallmark gene sets for each expression module shown in C. **H.** River plots showing the proportion of KLF5, AP1 and RUNX2 bound sites linked to the expression of their respective direct gene targets in the closed chromatin of senescent CD8+ T cells (represented as the color-coded modules as described in C).

**Figure S7.**
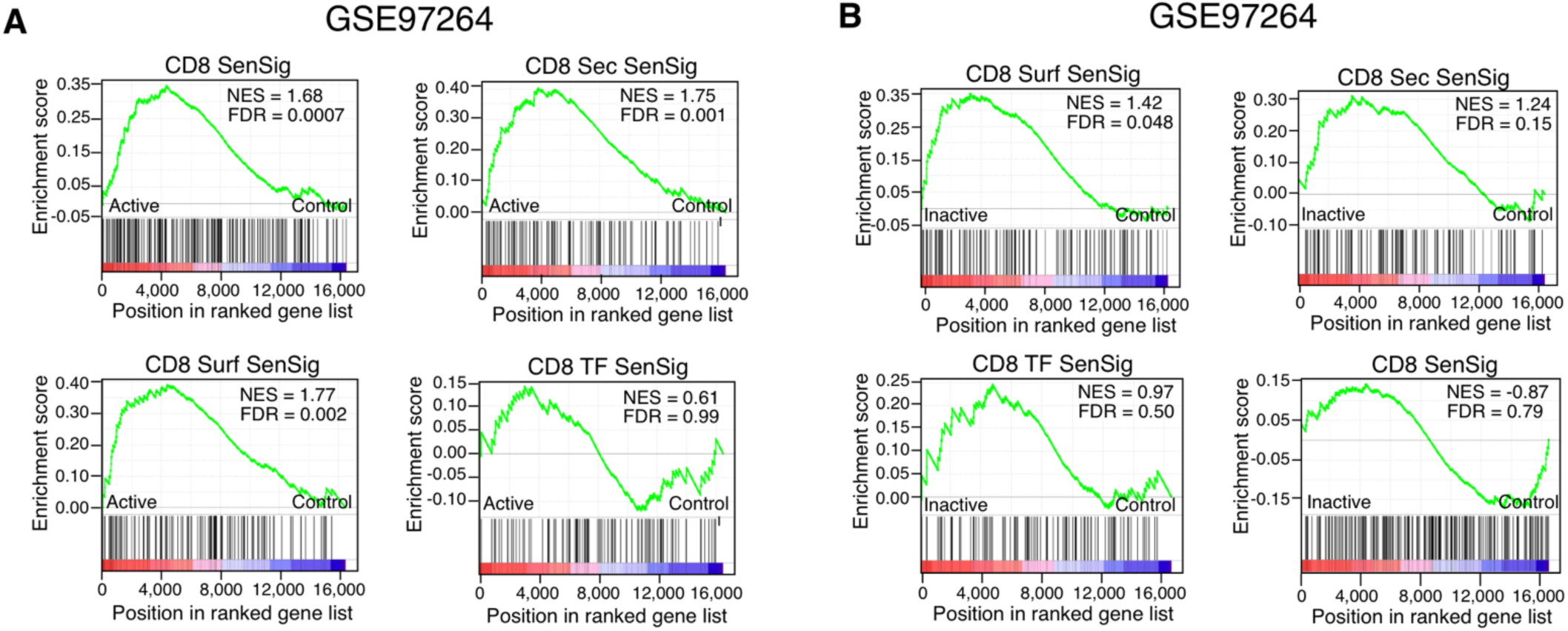
Predictive potential of CD8+ T cell senescence gene signatures of active systemic lupus erythematosus (SLE). A, B. Gene set enrichment analysis (GSEA) showing normalized enrichment score (NES) plots and FDR values for overall senescence (SenSig), senescence-specific surface-predicted (Sec SenSig), secreted-predicted (Surf SenSig) and TF (TF SenSig) gene signatures in an analysis of transcriptomes of CD8+ T cells from control donors and patients with active (A) and inactive (B) SLE from dataset GSE97264.

